# Mechanosensing through talin 1 contributes to tissue mechanical homeostasis

**DOI:** 10.1101/2023.09.03.556084

**Authors:** Manasa V.L. Chanduri, Abhishek Kumar, Dar Weiss, Nir Emuna, Igor Barsukov, Muisi Shi, Keiichiro Tanaka, Xinzhe Wang, Amit Datye, Jean Kanyo, Florine Collin, TuKiet Lam, Udo D. Schwarz, Suxia Bai, Timothy Nottoli, Benjamin T Goult, Jay D. Humphrey, Martin A Schwartz

## Abstract

It is widely believed that tissue mechanical properties, determined mainly by the extracellular matrix (ECM), are actively maintained. However, despite its broad importance to biology and medicine, tissue mechanical homeostasis is poorly understood. To explore this hypothesis, we developed mutations in the mechanosensitive protein talin1 that alter cellular sensing of ECM stiffness. Mutation of a novel mechanosensitive site between talin1 rod domain helix bundles 1 and 2 (R1 and R2) shifted cellular stiffness sensing curves, enabling cells to spread and exert tension on compliant substrates. Opening of the R1-R2 interface promotes binding of the ARP2/3 complex subunit ARPC5L, which mediates the altered stiffness sensing. Ascending aortas from mice bearing these mutations show increased compliance, less fibrillar collagen, and rupture at lower pressure. Together, these results demonstrate that cellular stiffness sensing regulates ECM mechanical properties. These data thus directly support the mechanical homeostasis hypothesis and identify a novel mechanosensitive interaction within talin that contributes to this mechanism.

## Introduction

The extracellular matrix (ECM) is the primary determinant of structural and mechanical integrity of most tissues. Maintaining ECM properties is essential to normal tissue function such that changes to ECM mechanics are central to pathological remodeling (*1*) in diverse diseases including cancer, coronary artery disease and kidney failure among others (*2, 3*). This connection between ECM organization and mechanics is especially crucial in the cardiovascular system. Heart failure is closely associated with fibrosis of cardiac tissue while stiffening of central arteries is a significant causal factor in cardiovascular, neurovascular and renovascular disease (*4*). Abnormal ECM remodeling plays a central role in aneurysms, the abnormal expansion of large arteries that predisposes to often-fatal dissection and rupture (*5*). Ascending thoracic aortic aneurysms (TAAs) in particular are principally caused by mutations in genes that code for ECM proteins such as fibrillin-1 (*6, 7*), contractile proteins such as smooth muscle α-actin (*7, 8*) and myosin (*9*), and upstream regulators of their expression and function such as TGFβ receptors (*10*) and myosin light chain kinase (*11*).

The compositional and organizational features of ECM that dictate mechanical properties are relatively constant over many decades of human life despite the much faster turnover of cells and most ECM components (*1, 12*). This observation has given rise to the concept of tissue mechanical homeostasis (*4*), whereby cells sense mechanical loads and properties of the ECM and adjust rates of matrix synthesis, assembly and degradation to maintain healthy tissue. Failure of these mechanisms is central in tissue fibrosis and aneurysms. The best evidence for homeostasis probably comes from studies of arteries, where changes in blood pressure that change circumferential stress leads to wall thickening that restores wall stress to nearly the original values (*13*). We recently reported a key role for microRNAs (miRNAs) in vascular endothelial cells, such that stiff environments upregulate a network of miRNAs that limit expression of contractile, adhesion and ECM proteins that promote tissue stiffening (*14*). Nevertheless, despite its conceptual appeal, experimental evidence supporting the concept of tissue mechanical homeostasis and mechanistic insights are sparse.

ECM stiffness is sensed by cells via transmembrane integrins and associated cytoskeletal linkers, adapters and signaling proteins, which can modulate intracellular biochemical pathways to elicit homeostatic responses (stiffness-sensing) (*15*). The load-bearing integrin-cytoskeleton protein talin 1 (hereafter talin) is a mechanosensitive linker that unfolds and refolds in response to tension (*16–19*). Changes in ECM or substrate stiffness determine the force applied to talin through modulation of contractile forces exerted by the cell (*18*), consistent with its role in stiffness sensing.

Talin contains a long, C-terminal rod domain consisting of sequential helix bundles, termed R1 through R13, that exhibit spring-like behavior, unfolding under tension and refolding when tension diminishes (*16, 17*). These domains function as bidirectional switches, wherein specific molecular interactions or signaling pathways are turned on or off by tension (*20*). Most prominently, vinculin binds to specific sites within many of the helix bundles (*21*), which reinforces the connection to actin under applied tension (*19*).

Deletion of talin or segments of the rod domain alters cellular mechanoresponses (*22, 23*). However, these large perturbations also alter integrin-mediated adhesion and signaling more generally, thus, are poorly suited to studying mechanical homeostasis *in vivo*. Here, we sought to test the contribution of cellular stiffness sensing to tissue mechanical homeostasis by developing mutations in talin that quantitatively shift stiffness sensing parameters without otherwise perturbing function. We generated mutations that mildly destabilize individual helix bundles. These experiments led to identification of the R1R2 interface as a novel mechanosensitive site that regulates cellular stiffness-sensing through its interaction with ARPC5L, an alternatively spliced component of the ARP2/3 complex. Mice bearing this mutation show altered ECM and mechanical properties in the ascending aorta. Together, these results identify a novel talin-dependent mechanotransduction mechanism and demonstrate that cellular stiffness sensing is critical for tissue mechanical homeostasis.

## Results

### Talin rod mutations

To investigate talin’s role in cellular mechanosensing and tissue homeostasis, we introduced mutations into individual rod domain helix bundles to decrease their stability though retaining their conformation at 37°C. Talin helix bundles are held together mainly by hydrophobic interactions within the core, thus, key hydrophobic amino acids were converted to more hydrophilic residues in the R1-R2 (L638D, L716T, A718D,V722R), R3 (I805T) and R10 (L1923A) domains to generate three destabilized talin rod-domain mutants (Fig. 1A). These domains were chosen because they contain well documented vinculin binding sites and are outside the actin, integrin binding and autoregulatory regions, thus, should not otherwise perturb the integrin-actin mechanical linkage. We performed circular dichroism to determine the thermodynamics of unfolding of the mutants in comparison to the wild-type domains. The indicated mutations led to moderate leftward shifts of the melting curves for those domains (Fig. 1B-1C, Fig S1A), though remained folded at 37°C. These mutations thus partially destabilize those domains and should lower the force required for unfolding.

**Figure 1:**
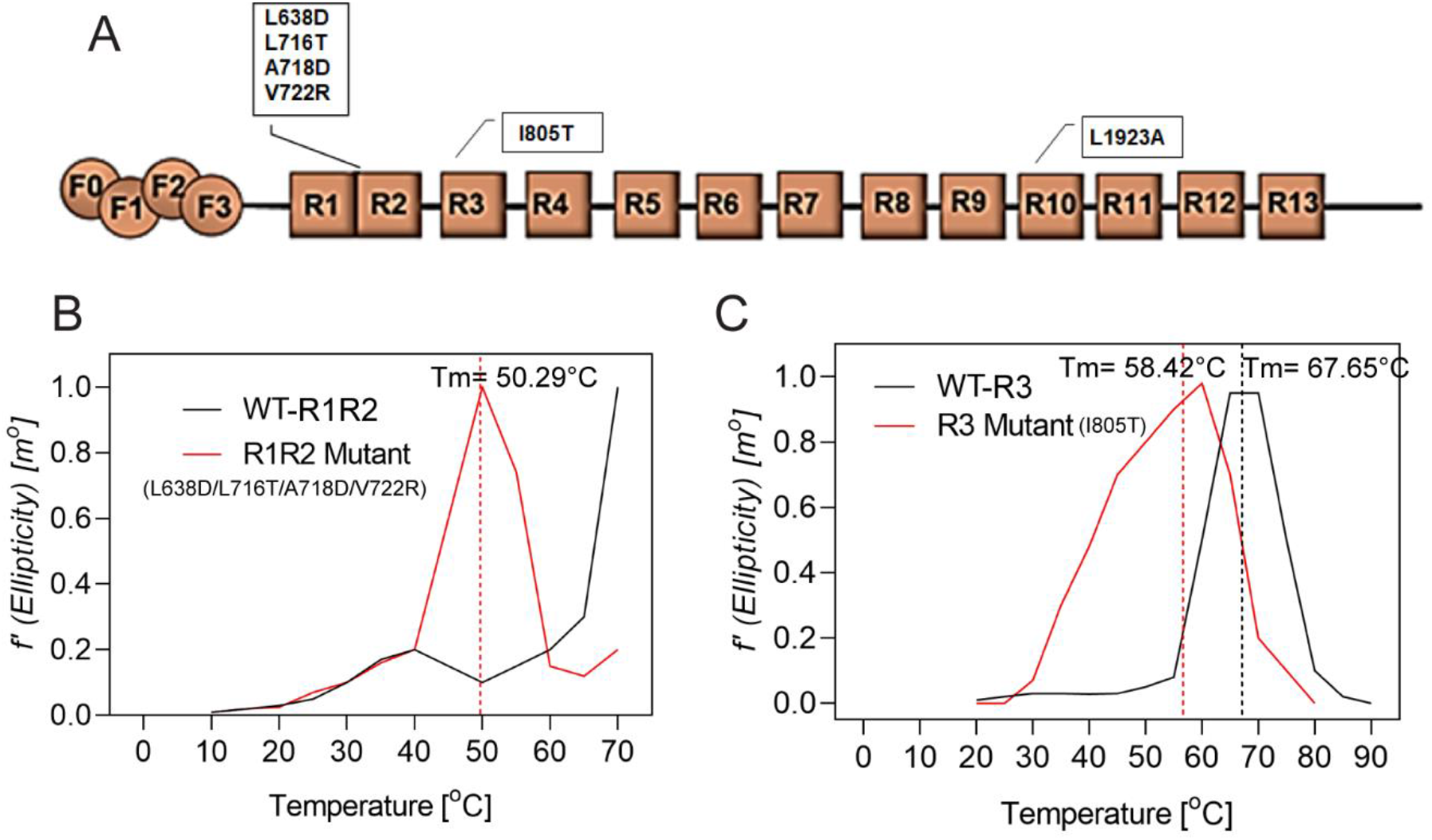
Mutations in rod-domains alter its stability. A) Domain organization of talin showing destabilizing mutations introduced into the R1R2, R3 and R10 domains. B-C) CD spectra showing molar ellipticity at a wavelength of 220 nm vs temperature in R1R2 (B) and R3 (C) mutant fragments compared to WT fragments. Dotted lines indicate the melting temperatures.

### Focal adhesion morphology and fibronectin deposition

These mutations were introduced into full-length talin containing the tension-sensor module to allow measurement of force across the molecule (*18*). Constructs were expressed in *Tln1*^−/−^ mouse embryonic fibroblasts (MEFs) at close to endogenous levels (Fig.S1B). All mutants localized to focal adhesions as expected (Fig. 2A). Cells expressing the R1R2 mutant had larger adhesions (Fig. 2A, B) compared to wild-type (WT) talin, whereas R3 and R10 mutants were similar to WT. The R1R2 mutant also showed increased recruitment to focal adhesions (Fig. 2C). Assessment of protein turnover within focal adhesions by fluorescence recovery after photobleaching (FRAP) showed no differences between the mutants and WT (Fig. 2D and Fig. S2A). Moreover, phosphorylation of focal adhesion kinase (FAK), and myosin light chain (MLC) were also unaffected, suggesting no change in focal adhesion signaling under these conditions (Fig. S2B-E).

**Figure 2:**
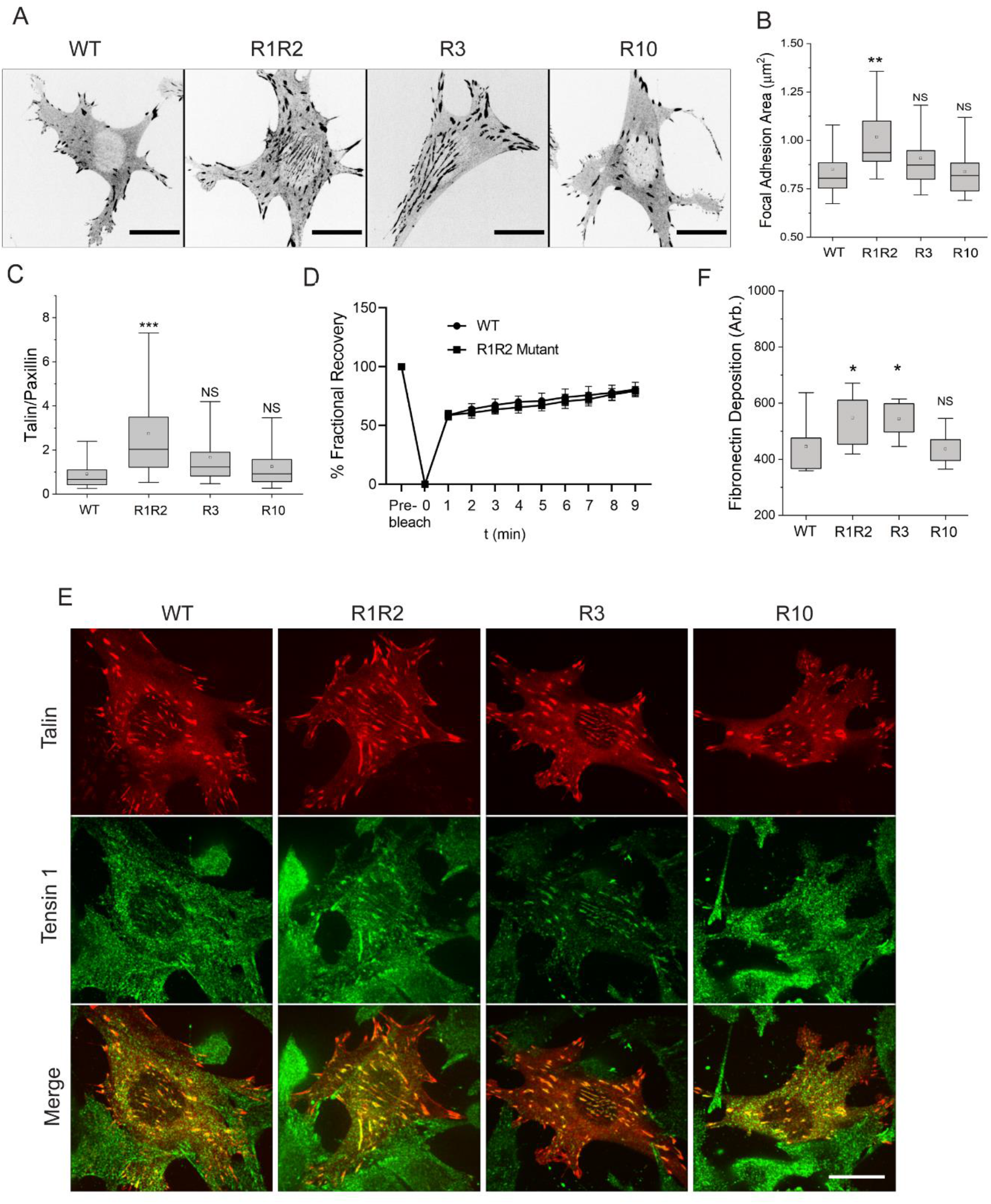
Effects of rod-domain mutations on focal adhesions. A) Representative intensity-inverted images showing focal adhesions in *Tln1^−/−^* MEFs expressing wild-type (WT) and rod-domain mutants plated for 3 h on a fibronectin-coated glass bottom dish. Scale bar, 20 µm. B) Quantification of focal adhesion area in A. N=25 cells for each sample. Boxes represent 25-75 percentile, whiskers represent 10-90 percentile, line is the median and dot represent the mean. Statistics were analyzed by one-way ANOVA, **, p<0.001, NS-Not significant. C) Quantified ratio of talin to paxillin within focal adhesions. N=25 cells in each box plot. Box represents 25-75 percentile, whiskers represent 10-90 percentile, line is the median and dot is the mean. Statistics were analyzed by one-way ANOVA, **, p<0.001, NS-Not significant. D) FRAP time course for WT and R1R2 mutant talin within individual focal adhesions. Data are means of % signal relative to starting values ± SEM. N = 10 cells per sample. E) Quantification of mean fibronectin intensity per cell in *Tln1^−/−^* MEFs expressing WT or mutant talin forms. N=15 fields of views from 3 experiments. Box represents 25-75 percentile, whiskers represent 10-90 percentile, line is the median and dot represent mean. Statistics were analyzed by one-way ANOVA, *, p<0.05, NS-Not significant. F) Representative immunofluorescence images of tensin (green) in *Tln1^−/−^* MEFs expressing wild-type (WT) and rod-domain mutants (red) plated on glass for 48 h.

Most of the cells expressing the R1R2 mutant showed large, elongated central adhesions (Fig. 2A) suggestive of fibrillar adhesions (*24*). We therefore stained for the fibrillar adhesion marker tensin-1. R1R2 mutant cells showed increased tensin-1 within the central adhesions (Fig. 2E and Fig. S3A). No changes in tensin-1 staining were observed with other mutants. Since fibrillar adhesions are sites of fibronectin deposition and matrix assembly (*24*)), cells were stained for fibronectin. Cells expressing the R1R2 mutant showed increased fibronectin fibrils (Fig. 2B and Fig. S3B) and increased fibronectin deposition by Western blotting (Supplementary Fig. S3 C, D). The results indicate that R1R2 regulates adhesion size and promotes formation of fibrillar adhesions with increased fibronectin fibrillogenesis.

### R1R2 mutant force transmission and substrate stiffness-sensing

Unfolding of R1R2 domains is expected to recruit more vinculin, strengthening the link to F-actin and increasing force transmission. We therefore assessed mechanical loading using our previously described fluorescence resonance energy transfer (FRET) sensor (*18*), for which decreased FRET indicates higher mechanical loading. When plated on glass coverslips, the talin R1R2 mutant showed decreased FRET (Fig. 3A), indicating higher tension, while the other mutants were comparable to WT.

**Figure 3:**
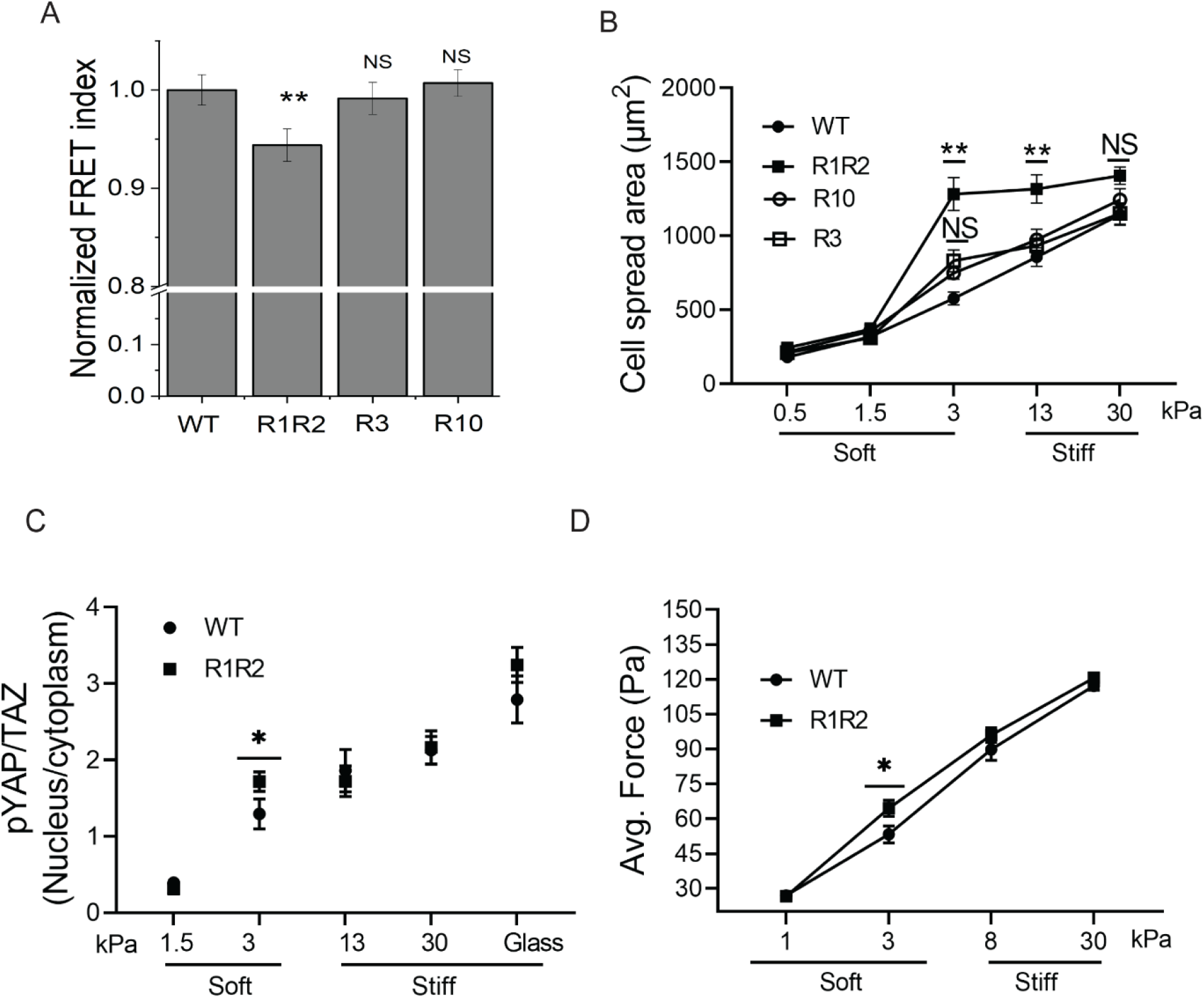
Effect of talin mutation on tension and stiffness-sensing. A) Normalized FRET index of WT and mutant talin in focal adhesions. Values are means ± SEM of N= 72-77 cells from three independent experiments. Statistics were analyzed by one-way ANOVA. **, p<0.001; NS- not significant. B) Quantification of spread area of cells expressing WT and talin mutants on polyacrylamide gels of varying stiffness. Values are means ± SEM of N= 52-78 cells. Statistics analyzed by one-way ANOVA, **, p<0.001; NS- not significant. C) Quantification of YAP localization in cells expressing WT and R1R2 mutant talin on soft substrates. Values are mean ± SEM of N= 25 cells per sample. Statistics analyzed by one-way ANOVA, *, p<0.05; NS- not significant. D) Average stress (force per unit area) in WT and R1R2 mutant-expressing cells on 3 kPa and 30 kPa substrates. Values are mean ± SEM of N= 25 cells. Statistics analyzed by one-way ANOVA. *, p<0.05; NS- not significant.

Cell sensing of substrate stiffness has been linked to talin (*18, 22*) and domain opening could be expected to alter this behavior. Cells expressing WT vs mutant talin constructs were therefore plated on fibronectin-coated polyacrylamide hydrogels with varying stiffness and cell morphology assessed. Cells containing the R3 and R10 mutant showed the expected increase in cell area with increasing substrate stiffness comparable to WT talin (Fig. 3B). By contrast, the R1R2 mutant induced a distinct leftward shift in the curve, such that these cells were fully spread on a compliant/soft 3 kPa substrate, whereas control cells required 30 kPa (stiff substrate) to achieve a similar spread area (Fig.3B and Fig. S4A). To determine whether this shift led to changes in signaling, we examined YAP/TAZ, transcription factors that show stiffness-dependent nuclear translocation and control subsequent cellular responses (*25*). Talin R1R2 mutant cells also increased Yap/Taz nucleus-to-cytoplasmic ratio on the 3 kPa substrate (Fig. 3C and Fig. S4B). We next examined traction stress in cells on 3 vs 30 kPa substrates. Cells increase forces exerted on substrates as a function of spread area (*26*). However, expressing these tractions as stress (force/unit area) corrects for the contribution from higher spreading, allowing assessment of area-independent effects (Fig. 3D and Fig. S4C). Traction stress in R1R2 mutant cells also showed a leftward shift, with higher stress on surfaces of intermediate stiffness. Together, these results provide strong evidence that the R1R2 mutant alters cell substrate stiffness-sensing.

### Vinculin-dependent and independent effects

Helix bundle unfolding promotes vinculin binding (*16, 27*) and increases the tension on talin (*18, 28*). To test the requirement for vinculin in these effects, WT and R1R2 mutant talin were expressed in talin and vinculin double knockout cells (Fig. S5A) and the above parameters assessed in cells plated on glass coverslips. Loss of vinculin abolished the difference in tension on talin between the WT and R1R2 mutant in cells in cells lacking both endogenous talin and vinculin (Fig. 4A). However, there was only a partial rescue of focal adhesion size. Importantly, when examined on polyacrylamide gels, while maximal spread area was diminished upon vinculin depletion, the shift in the stiffness sensing curve with the R1R2 mutant was preserved, with spreading still maximal on 3 kPa substrates (Fig. 4B-4D). These results reveal both vinculin-dependent (talin tension) and vinculin-independent (stiffness sensing) effects, with focal adhesion size showing components of both. These findings suggest the existence of tension-dependent effectors of R1R2 unfolding other than vinculin.

**Figure 4:**
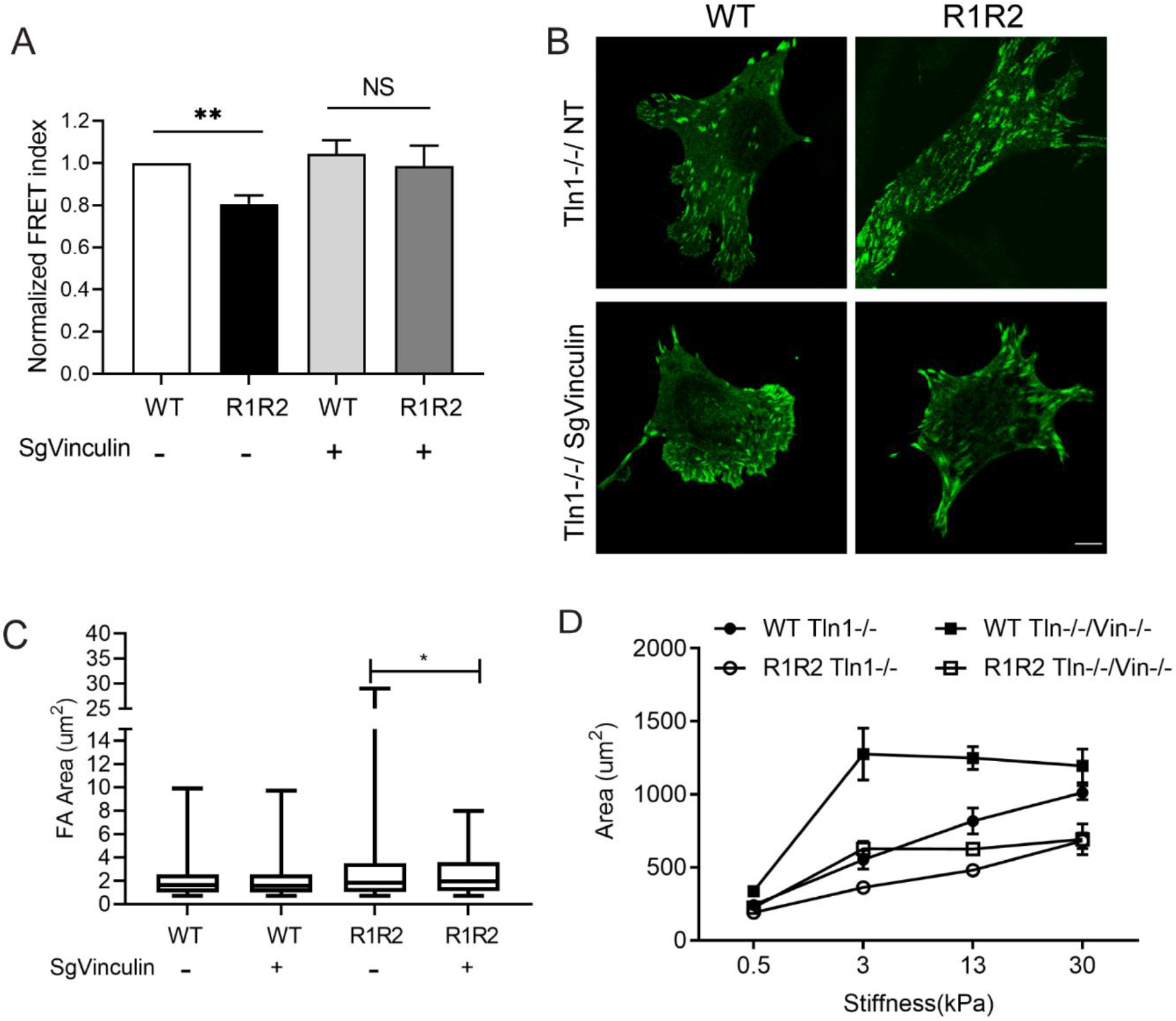
Effect of vinculin depletion on R1R2 mutant phenotypes. A) Normalized FRET index of WT and R1R2 mutant talin expressed in *Tln1^−/−^* MEFS after vinculin CRISPR KO. Values are means ± SEM, n= 25 cells. Statistics analyzed by one-way ANOVA. NS-Not significant; **, p<0.01. B) Representative images of WT- and R1R2-expressing *Tln1^−/−^* MEFS with or without vinculin KO. Scale bar, 5 µm. C) Focal adhesion area in *Tln1^−/−^* MEFs with or without vinculin KO. Values are means ± SEM of N=52-102 cells. D) Quantification of cell area in WT and R1R2 expressing cells with and without vinculin KO. Values are means ± SEM of N= 32-54 cells.

### Critical role for the R1-R2 interface

The R1R2 domain are uniquely organized, with R1 and R2 making a side-to-side contact in addition to the end-to-end contact that occurs in all domains (*27*). The above effects might be due to opening of either the R1-R2 interface or the R2 helix bundle core. To address this question, we developed four R1R2 domain mutants: two mutants (L716T/L682T/ V747T and L682T/V747T) that target the R2 internal core interactions (Fig. 5A and Fig S5B) and two (L638D/V722R and L638D/A718D/V722R) that target the R1R2 interface but leave the R2 core unperturbed (Fig. S5B). These mutations were introduced into full-length talin and expressed in *Tln1^−/−^* MEFs. When assayed for spreading on 3 vs 30 kPa substrates, only the R1R2 interface mutations increased spreading on soft substrate (Fig. 5B). The interface mutants also had larger adhesions (Fig. 5C). However, the change in tension on talin, indicated by FRET index, was not seen with either the core or interface mutations alone (Fig. 5D), indicating that this effect required both. As vinculin binding requires exposure of the helix bundle core, these results are consistent with findings from Fig. 4 that effects on cellular stiffness sensing and focal adhesion size occur through an effector other than vinculin.

**Figure 5:**
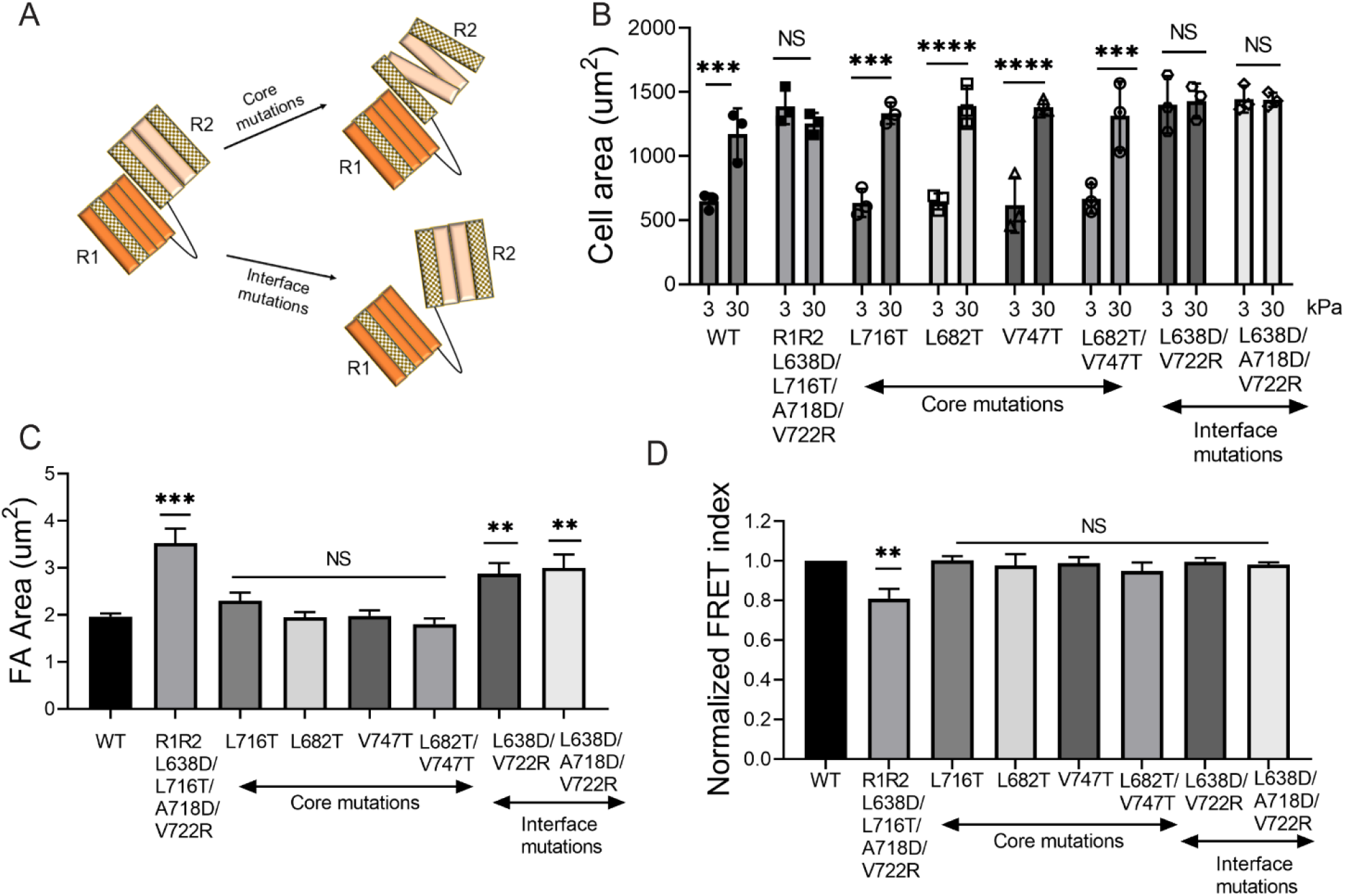
Effects of R2 core vs R1-R2 interface interactions. A) Schematic of R2 core vs R1R2 interface mutations. B) Cell spread area of WT, R1R2 core and interface talin mutants on 3 kPa and 30 kPa substrates. Values are means ± SEM of N= 3 experiments. Statistics analyzed by one-way ANOVA. NS-Not significant; ***, p<0.001; ****, p<0.0001 C) Focal adhesion area of WT, R1R2 core and interface mutant expressing *Tln1^−/−^* MEFs. Values are means ± SEM of N=.30 cells. Statistics analyzed by one-way ANOVA. NS-Not significant; **, p<0.01; ***, p<0.001

### ARPC5L binds the R1R2 interface

To identify effectors that bind the R1R2 interface, we reasoned that proteins that bind free R2 domain but not the R1R2 dimer would be potential effectors that interacted only after tension-dependent disruption of R1-R2 binding. We therefore performed pull-downs from cell lysates expressing GFP-tagged R2 and GFP-tagged R1R2 domains, to identify binding partners by mass spectrometry (Table S1). More than 30 proteins preferentially bound free R2 domain. These included multiple cytoskeletal proteins that are potential effectors of cellular mechanosensing (Table S2). To narrow the search, we first assessed whether the effectors worked through F-actin or microtubules. Thus, stiffness-dependent cell spreading was assayed in the presence of the ARP2/3 inhibitor CK666, the formin inhibitor SMIFH2 and the microtubule inhibitor nocodazole. The R1R2 stiffness-sensing phenotype was reversed by CK666 but not by the other inhibitors (Fig. 6A). CK666 also partially reversed the large adhesion size in cells expressing the R1R2 mutant talin but had no significant effect on cells expressing WT talin (Fig. 6B-6C).

**Figure 6:**
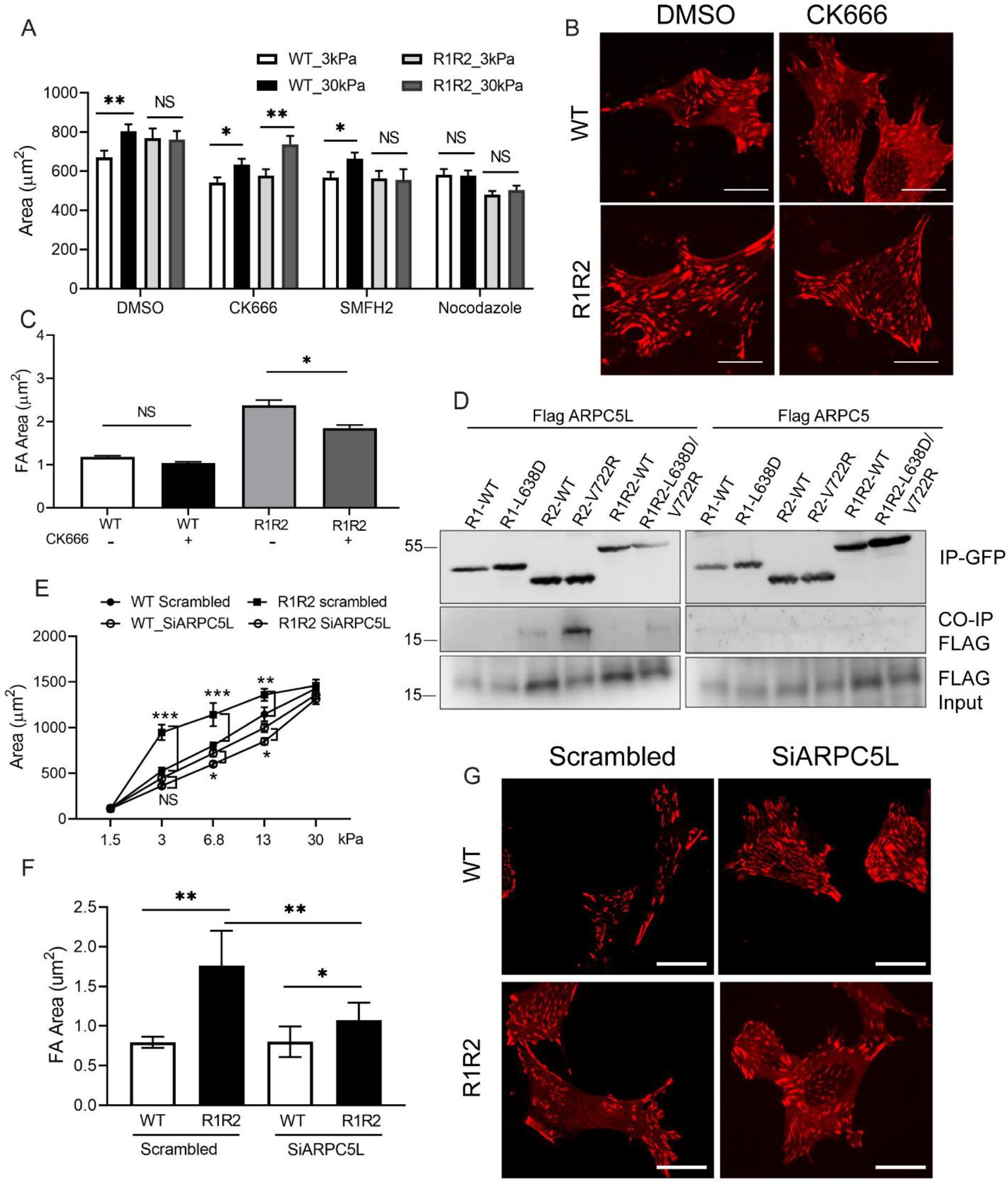
ARPC5L mediates talin R1R2-mediated stiffness-sensing. A) Cell area in WT and R1R2 expressing cells on 3 and 30 kPa stiffness substrates in the presence of DMSO, ARP2/3 inhibitor (CK666), formin inhibitor (SMIFH2) and nocodazole (microtubule polymerization inhibitor). Values are means ± SEM of N=25-50 cells. Statistics analyzed by one-way ANOVA. NS-Not significant; *, p<0.05; **, p<0.01. B) Representative images showing focal adhesions with and without CK666 in cells expressing WT and R1R2 mutant. Scale bar, 10 µm. C) Quantified focal adhesion area from (B). Values are means ± SEM of N= 25 cells. Statistics analyzed by one-way ANOVA. NS-Not significant; *, p<0.05. D) Representative blots showing co-immunoprecipitation of Flag-tagged ARPC5L and Flag-tagged ARPC5 with the indicated GFP-tagged talin constructs. E) Quantified cell spread area in WT and R1R2 expressing cells with and without ARPC5L depletion on hydrogels of varying stiffness. Values are means ± SEM of N=25-50 cells. Statistics analyzed by one-way ANOVA. NS-Not significant; *, p<0.05; **, p<0.01; ***, p<0.001. F) Quantification and G) representative images of WT and R1R2 mutant expressing cells with and without ARPC5L depletion. Scale bar, 10 µm. Values are means ± SEM of N= 45-72 cells. Statistics analyzed by one-way ANOVA. *, p<0.05; **, p<0.01.

These results led us to investigate ARPC5L, one of the proteins that specifically bound the R2 domain in the proteomic assay (Table S2). ARPC5L is a component of the ARP2/3 complex that nucleates F-actin assembly. To confirm the interaction, GFP-tagged version of WT and mutant constructs of R1R2, R1 and R2 domains were expressed in cells together with Flag-tagged ARPC5L or ARPC5 (a functional isoform of ARPC5L). Lysates were immunoprecipitated with GFP nanobody beads and analyzed by Western blotting. ARPC5L showed little association with WT R1R2 or WT R1 but significantly more with WT R2. ARPC5L showed higher association with the mutant R1R2 and higher still with the mutant R2 domain. ARPC5 showed no binding to any of the talin constructs. These results support the notion that ARPC5L binding requires separation of the R1 and R2 domain and is specific to this isoform (Fig. 6D). Further, computational modeling of the R2 domain showed strong binding with ARPC5L including when it is part of the ARP2/3 complex (Fig.S6). Modeling also identified the ArpC5L insert region that is absent from ArpC5, consistent with the observed binding specificity. To functionally test this hypothesis, we depleted ARPC5L in cells expressing the R1R2 interface mutant. ARPC5L knockdown reversed the altered stiffness-sensing and increased adhesion size in the R1R2 mutant (Fig. 6E-6G). ARPC5L depletion in cells expressing WT talin also slightly but significantly decreased cell spreading on substrates of 6.8-13 kPa. Together, these results identify ARPC5L as a novel effector of talin mechanosensitivity.

### Mice with R1R2 interface mutations

To understand effects of altered stiffness-sensing at the organismal level, the interface mutations (L638D and V722R) were introduced in C57BL/6 mice using CRISPR-Cas9 mutagenesis (Fig. 7A, Fig S6A). The mutant mice (addressed henceforth as R1R2 mutant mice) were viable (Fig. 7B) and fertile (Fig. S6B). Younger mice showed slightly diminished weight, but this difference disappeared by 4 weeks (Fig. 7C). As mutations in ECM and contractile proteins play a critical role in mechanical homeostasis of the ascending aorta (*4*), we carried out a mechanical analysis of this tissue. Atomic force microscopy measurements on the ascending aorta from mice at P24 revealed a lower compressive elastic modulus in the mutant tissue compared to WT littermates (Fig 7D). Burst pressure, tensile wall stiffness and stress under biaxial loading at physiological levels were also lower in ascending aortas from R1R2 mutant mice (Fig. 7F-7G). As expected from the reduced stiffness, circumferential stretch using typical values for *in vivo* diastolic and systolic pressures was greater in the R1R2 mutant (Fig. 7H). However, other mechanical properties that we assessed were unchanged (Fig. 7I-7J). To our surprise, comparable analysis of the descending aorta showed no difference (Fig. S7). These results suggest that altering cellular stiffness sensing with the R1R2 mutation leads to reduced tissue stiffness specifically in the ascending aorta.

**Figure 7:**
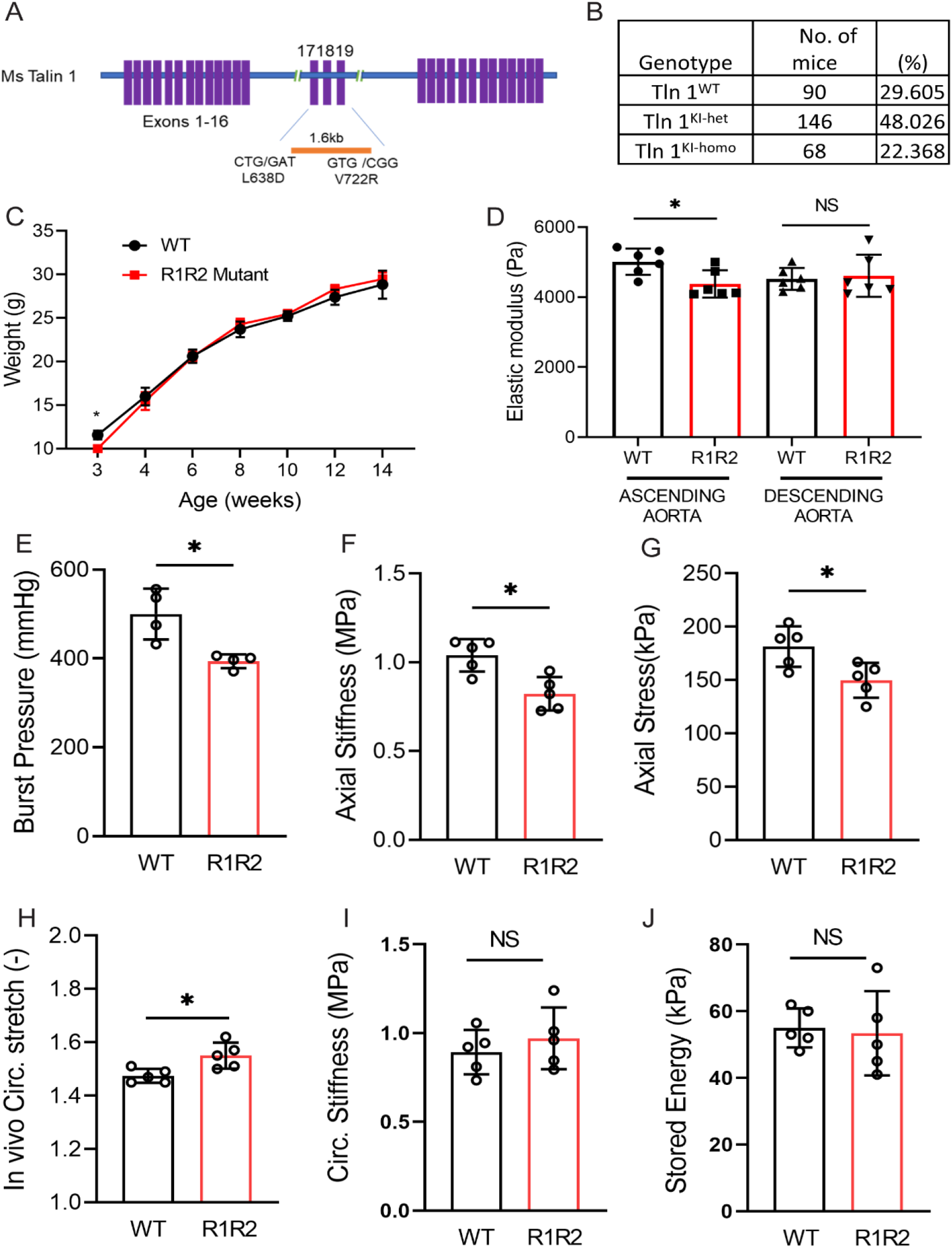
R1R2 mutant knock-in mice. A) Schematic showing the insertion of mutations L638D and V722R in exon 17 and exon 19 using a 1.6 kb template. B) Genotype of progeny from mating heterozygous *Tln1*^L638D-V722R/+^ mice (*Tln1*^KI-het^). Chi-square analysis revealed no significant difference. Chi-squared equals 3.658 with 2 degrees of freedom. The two-tailed P value equals 0.1606 and is not statistically significant. C) Body weight of WT and R1R2 mutant knock-in mice from 3 to 14 weeks of age. Values are means ± SEM. Statistics analyzed by unpaired t-test. *, p<0.05. D) Compressive radial stiffness of the ascending aorta of P24 WT and R1R2 mutant mice measured by atomic force microscopy applied from the adventitial surface. Values are means ± SEM of N= 5 mice. Statistics analyzed by unpaired t-test. NS-Not significant; *, p<0.05. E-J) *Ex* vivo biomechanical measurements of ascending aorta of P24 WT and R1R2 mutant mice showing changes in burst pressure (E), tensile axial stiffness (F), tensile axial stress (G), *in vivo* extensional circumferential stretch (H), tensile circumferential stiffness (I) and stored energy density (J).

ECM is the main determinant of the mechanical properties of the arterial wall (*4, 29*). A change in stiffness thus points toward altered ECM. Histological analysis of fibrillar collagen by picrosirius red and Movat pentachrome revealed reduced staining in the R1R2 mutant aorta in both the adventitial and medial layers (Fig. 8A-8C). Collagen III was also lower in the mutant tissues (Fig. S8). Taken together, R1R2 mutation with altered stiffness-sensing *in cellulo* results in reduced ECM deposition and decreased material stiffness *in vivo* in the mouse ascending aorta.

**Figure 8:**
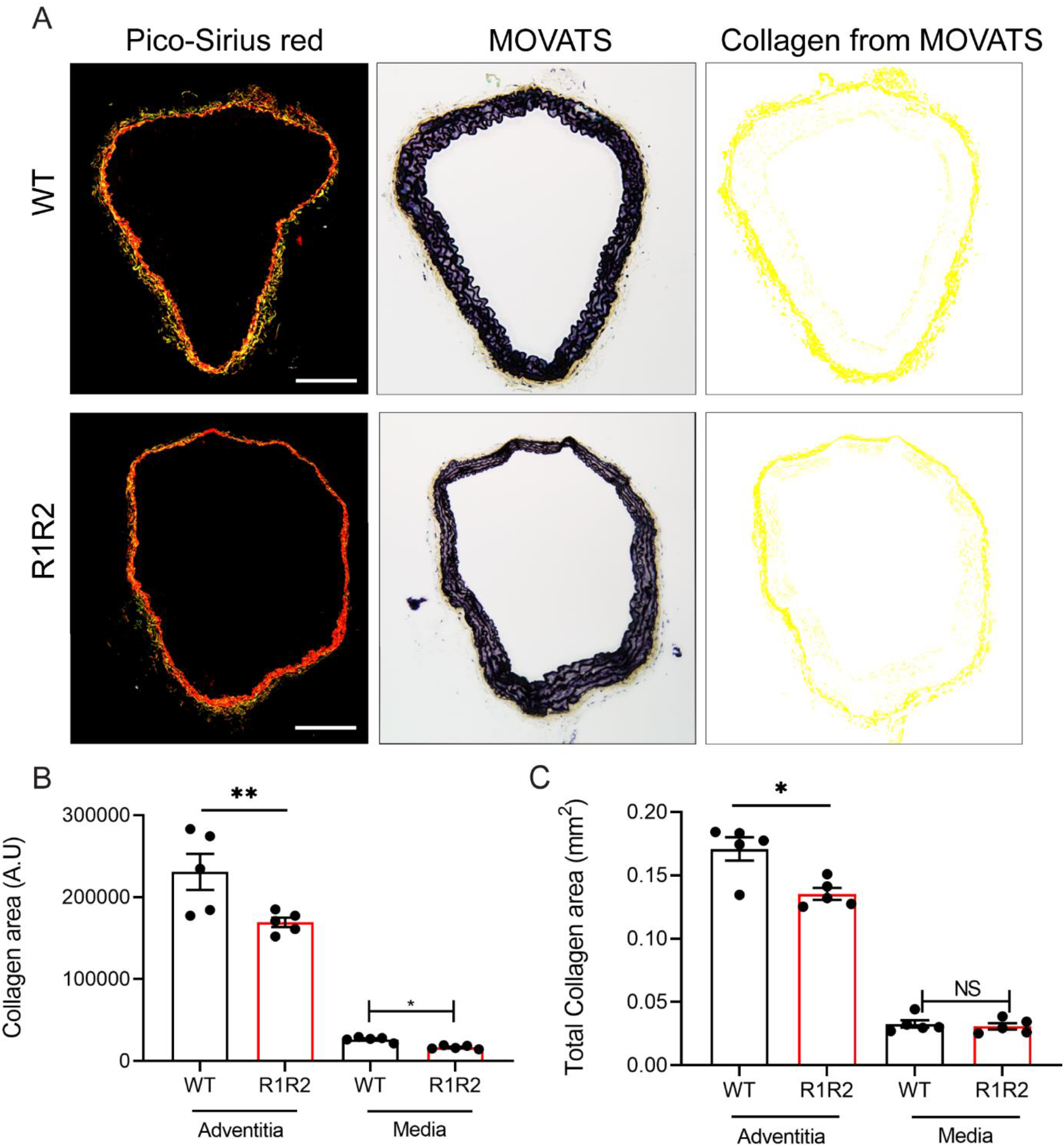
ECM in R1R2 mutant aorta. A) Picrosirius red staining (left panel) and Total Movat pentachrome staining (middle panel). Collagen content in Movat stains extracted by custom MATLAB software (*66*) (right panel) for ascending aortas from WT and R1R2 mutant mice at P24. Scale bar, 100 µm. B) Quantification of total collagen fibers identified by picrosirius red staining in the adventitia and media. Values are means ± SEM of N= 5. Statistics analyzed by unpaired t-test. NS- Not significant; *, p<0.05; **, p<0.01 C) Quantification of total collagen area within adventitia and media of ascending aorta at P24 by Movat staining. Values are means ± SEM of N= 5. Statistics analyzed by unpaired t-test. NS- Not significant; *, p<0.05.

## Discussion

The goal of this study was to explore the connection between cellular stiffness sensing and tissue mechanical homeostasis, focusing on ECM as the main determinant of tissue stiffness. We approached this challenge by developing talin mutants in which stability of specific rod domain helix bundles was reduced, which should allow opening under lower tension. We found that mutation of the R1-R2 interface had the strongest effect, increasing cell spreading and traction forces on surfaces of 2-3-fold lower stiffness. When introduced into mice, these mutations resulted in reduced stiffness and strength of the ascending aorta that correlated with reduced fibrillar collagen. Thus, when cells (incorrectly) sense the ECM as too stiff, they decrease collagen assembly to reduce tissue stiffness. This outcome provides direct support for the concept that cellular stiffness sensing governs tissue mechanical homeostasis.

A second key finding is identification of the R1-R2 interface as a force-dependent site. R2 that is released from the R1 interaction but still in its folded conformation can bind ARPC5L to promote cell spreading. Single molecule experiments showed that an R1-R3 construct unfolded in three distinct steps with large scale movements attributable to R3 at ∼5 pN, R2 ∼13 pN and R1 at ∼22 pN respectively (*30*). Additionally, a smaller movement was observed at ∼10 pN, prior to full opening of R1 or R2 (Figs. 3a and e in (*30*)). While this point remains to be fully investigated, the notion that R1-R2 separation occurs at a relatively early stage is consistent with available data.

The original R1R2 mutant that contained both interface and core mutations also showed increased formation of fibrillar adhesions and increased tension on talin. Both effects were diminished by vinculin depletion, consistent with increased vinculin recruitment to this mutant. These effects were much diminished in talin constructs that contained interface mutations only.

ARPC5L is a component of the ARP2/3 complex (*31*) and an orthologue of ARPC5. ARPC5L shares 68% identity with ARPC5 and specifically differs by an insertion of six alanine residues in its N-terminus (Fig. S6A). The specific binding of ARPC5L implies that this insert is required for binding to talin R2. In fact, modeling of ARPC5L and talin R2 fragment revealed a specific interaction between the free R2 domain and the six alanine residues in ARPC5L (Fig. S6B), also seen with ArpC5L assembled in the ARP2/3 complex (Fig. S6C). ARPC5L (or ARP5L) was found in the focal adhesion proteome of blebbistatin-treated fibroblasts along with other ARP2/3 components (*32*). Previous studies identified interactions of the ARP2/3 complex with vinculin and FAK (*33–35*) that reportedly regulate cell migration and lamellipodia formation. During migration, interaction of FAK with the ARP2/3 complex was found to be important for haptosensing of ECM concentration but not for ECM stiffness-sensing (*36*). Melanoblasts derived from *Arp3* knockout mouse embryos also displayed decreased spreading on soft tissues and loss of cell-ECM contacts on stiffer tissues (*37*). Identification of a novel tension-sensitive interaction between talin and ARP2/3 thus strengthens the case for a key role of this actin regulator in cell adhesion.

Talin1 was identified as a potential causal gene in sporadic thoracic aortic aneurysms (*38*) and as a susceptibility gene for spontaneous coronary artery dissection (*39*). Talin1 was also reported to be downregulated in aortic dissections (*40*). While these disease connections remain to be confirmed by functional analyses and larger scale genetic studies, they lend support to the notion that talin plays a role in mechanical homeostasis of arteries. One surprising result was the specificity for the ascending relative to descending aorta. This effect could be due to the distinct developmental origins of vascular smooth muscle cells in these segments (*41*). Additionally, because each contraction of the heart pulls down on the ascending aorta, the ascending aorta undergoes both axial and circumferential strain during the cardiac cycle, whereas the descending aorta undergoes only circumferential strain (*42*). To what extent these developmental and biomechanical differences account for the difference in talin-dependent matrix assembly remains to be determined.

In our study, we observed a decrease in aortic stiffness in R1R2 mutant mice at P24. This age was chosen as a stage in development when elastin deposition in the aortic media is complete but pressure-induced collagen organization and compensatory mechanisms are emerging (*12*). Lower stiffness was associated with lower content of fibrillar collagen in both the adventitia and media, suggesting that more than one cell type is affected. How ECM structure and content will evolve with age is an ongoing question.

In summary, our work identified the R1R2 talin module as a critical stiffness regulator that trigger downstream stiffness-sensing responses through ARPC5L. The R1R2 mutant mice show subtle but significant differences in ECM and aortic mechanics in young mice. Important questions for future work include identifying pathways that link stiffness sensing to ECM gene expression, assembly and turnover; the contribution of substrate-stiffness sensing in mechanical homeostasis in other tissues; how this pathway participates in age-related artery stiffening; and how homeostatic regulation fails in diseases such as aneurysms and arteriosclerosis.

### Limitations of this study

We cannot exclude unintended consequences of mutations within the R1-R2 region, for example, alterations in binding to additional components. As the R1-R2 interaction contributes to the stability of both the R1 and R2 domains, these mutations may reduce the force necessary to unfold either one. Binding of ARPC5L to R2 is predicted to block the R1-R2 interaction and to stabilize R2, which may affect other processes. Recruitment of ARPC5L to talin may have effects on ARP2/3 functions in other regions of the cells. Mice with R1-R2 interface mutations were examined at a single age chosen to minimize adaptation to altered talin function, but compensatory or pathological mechanisms cannot be excluded.

## Experimental Design

### Cell culture

*Tln1^−/−^* MEFs (*43*) were cultured as previously described (*18*). Briefly, MEFs were grown in cultured in DMEM/F12 with 10% FBS, penicillin-streptomycin, 0.001% β-mercaptoethanol and 0.12% sodium bicarbonate. CRISPR-Cas9 deletion of vinculin in *Tln1^−/−^* MEFs used the vinculin specific guides RNAs (sgNT 5’-GCGAGGTATTCGGCTCCGCG-3’; sgVinculin 5’-388 GCCGTCAGCAACCTCGTCC-3’) and procedures as described (*44*). These cells were cultured in *Tln1^−/−^* MEFs media with 1 µg/mL puromycin. All cells were maintained in a humidified 37°C, 5% CO_2_ incubator.

### Plasmids and transfection

The R1R2, R3 and R10 mutant fragments as well as the R1R2 core and interface mutations were inserted in the previously described full-length talin-TS (*18*) by Gibson assembly. Rod-domain fragments for circular dichroism: His-tagged R1R2 WT, R1R2 (L638, L716T, A718D, V722R) mutant, R3 WT and mutant (I805T) and R10 WT and mutant (L1923A) as well as R1R2 interface WT and mutant fragments were inserted into Ptrc-His A vector and expressed in BL21 DE3 *E. Coli* and purified on Ni-NTA column as described (*45*). For co-immunoprecipitation, flag-tagged ARPC5L was generated from pLVX GFP-ARPC5L (*31*) and GFP-tagged R1R2 WT and mutant fragments as well as R1 and R2 WT and mutant fragments were inserted in pLPCX vector. All cells were seeded one day before transfection and transfected using Lipofectamine^TM^ 2000 (ThermoFisher Scientific) as per manufacturer’s instructions. Cells were used in experiments 36 h post-transfection.

### Circular Dichroism, FRET and FRAP

Circular dichroism was used to assess stability of talin rod domain constructs. Measurements were conducted using 100 µM of talin fragment (WT, R1R2, R3 or R10 mutants) dissolved in buffer containing 50 mM KCl and 10 mM MES, pH 6.1. Spectra were recorded on a Jasco J-1100 CD spectrometer for the temperature intervals 10-70 and 20-80°C for R1R2, R3 and R10, respectively, using a 1mm path length and from wavelength range 250 to 180 nm. Spectra were collected every 5°C, with the temperature equilibration delay of 5 min. The scan rate was 100 nm/min, 3 accumulations per sample were averaged to enhance sensitivity. Melting temperatures were evaluated from the maximum of the first derivative of the temperature dependence of the ellipticity at 220 nm.

FRAP experiments to assess talin turnover were performed on a Leica SP8 confocal microscope equipped with a FRAP module using a 63x, 1.4 NA oil objective. Cells were maintained at 37°C with humidity and CO_2_ control. Images were acquired using LASX software. Three prebleach images at 2-s intervals and then a laser pulse at 100% power of the 488-nm line were used to bleach focal adhesions. Time-lapse images were acquired every 20 s for 9 min. Images were corrected for photobleaching during image analysis and normalized FRAP curves were plotted (*18*).

FRET imaging and analysis to measure tension on talin WT and mutant forms was performed as described (*18*). Briefly, *Tln1^−/−^* MEFs transfected with WT and mutant rod-domain plasmids were plated for 6 h on fibronectin-coated glass-bottom dishes. Images were acquired using a 100×, 1.4 NA oil objective of a Nikon TiEclipse in the donor, acceptor and FRET channels and normalized FRET index was calculated using a customized MATLAB code.

### Focal adhesion morphology, Fibronectin deposition and stiffness-sensing assays

To assess \ focal adhesion morphology, cells transfected with GFP-tagged WT or mutant talin were seeded for 6 h on glass-bottom dishes coated with fibronectin at 10 µg/mL, then fixed with 4% paraformaldehyde. Cells were imaged using a 60x 1.4 NA oil objective on a Nikon Eclipse Ti microscope and focal adhesion areas analyzed by Image J. Co-localization of WT and mutant talin was performed by immunostaining with antibodies against phospho-paxillin (rabbit; 44-722G; ThermoFisher Scientific), vinculin (mouse, V9264, Sigma), phospho-myosin light chain 2 (ser19) (rabbit; 3671; Cell Signaling Technology). For fibrillar adhesion analysis, *Tln1^−/−^* MEFs expressing WT and R1R2 mutant talin were grown for 48 h on glass-bottom dishes, fixed with 4% paraformaldehyde and stained for tensin 1 (rabbit; SAB4200283; Sigma-Aldrich; 1:300). Tensin 1 intensity within central adhesions was calculated as previously described (*46*). For fibronectin deposition, cells were grown for 48 h on glass-bottom dishes, fixed with 4% paraformaldehyde and stained for fibronectin using rabbit fibronectin (rabbit; F3648; Sigma-Aldrich; 1:300). Cells were counterstained with phalloidin to obtain cell boundaries. The fibronectin signal under each transfection positive cell was quantified using Image J.

Fibronectin fibrillogenesis was assayed biochemically as described (*47*). Briefly, *Tln1^−/−^* MEFS expressing WT and R1R2 talin were grown on 10 cm dishes for 48 h and washed in ice-cold PBS. Cells were lysed in a solubilization buffer, pH 8.8 [deoxycholate (2%), Tris-HCl (25 mM), PMSF (2 mM), iodoacetic acid (2 mM), N-ethylmaleimide (2 mM) and EDTA (2 mM)]. The insoluble fraction was pelleted at 16,000g for 15 min. The pellet was solubilized in SDS (2%), Tris-HCl (25 mM), PMSF (2mM), iodoacetic acid (2 mM), N-ethylmaleimide (2 Mm), and EDTA (2 mM), pH 8.0. The proteins were mixed with 4x Laemlli buffer (Bio-Rad), denatured at 95°C for 10 min, resolved on 8% SDS-PAGE gel and analyzed by Western blotting. The blots were incubated with anti-fibronectin antibody (rabbit; F3648; Sigma-Aldrich; 1:1000) overnight at 4°C followed by incubation with HRP-conjugated anti-rabbit IgG (goat; PI-1000-1; Vector Labs; 1:5000) and proteins bands were visualized by chemiluminescence.

For stiffness sensing assays, hydrogel substrates of varying stiffness were generated as described (*48*), by varying the acrylamide-bisacrylamide ratios. Fibronectin was covalently linked to the gels pre-mixed with N-hydroxy succinimide (NHS) ester (1 mg/mL; 6066-82-6; Chemcruz) by overnight incubation at 4°C. The gels were then washed with PBS and equilibrated in cell culture media 1 h prior to seeding. Cells were plated on these gels for 6 h, followed by fixing with 4% paraformaldehyde and stained with rhodamine-phalloidin to image the cell body or with anti-YAP antibody (mouse; sc-101199, Santa Cruz Biotechnology; 1:300) to determine Yap nuclear localization.

### Traction force microscopy

Traction force microscopy to assess cellular forces was performed as described (*18*). A thin-layer of PDMS substrate on glass-bottom dishes was prepared by spin-coating. 3 kPa stiffness substrates were made by mixing a 1:1 ratio of silicone and curing agent (CY-52-276A and CY-52-276B; Dow Corning). 30 kPa stiffness substrates were prepared by mixing a base and curing agent from Sylgard 184 (Dow Corning) in a 40:1 ratio. The surface of the PDMS substrate was silanized using 3-aminopropyltriethoxy silane and a bead solution comprising borate buffer, pH 7.4 and 0.2 µm fluorescent beads (F-8807, Invitrogen, Molecular Probes) (1:2500 beads: buffer; v/v) and 0.1 mg/mL 1-ethyl-3-(3-dimethylaminopropyl) carbodiimide (EDC) was added for 2 h at room temperature. The surface was then coated with 10 µg/mL fibronectin overnight at 4°C. Cells were seeded and allowed to spread for 6 h. Images of the beads with cells and after treatment with 0.1% SDS were acquired, and traction forces were calculated using custom-written MATLAB code (*49, 50*)

### Mass spectrometry

Plasmids encoding GFP or GFP-tagged R1R2 or GFP-tagged R2 wild-type constructs were expressed in 50% confluent HEK293T cells in a 100 mm dish using Lipofectamine 2000 as per manufacturer’s protocol. 36 h after transfections, cells were lysed in buffer containing 25 mM Tris, pH 7.4, 1% NP-40, 150 mM NaCl, 0.1% BSA, and 1x protease and phosphatase inhibitor cocktail (ThermoFisher) at 4°C for 30 min. Proteins were immunoprecipitated using GFP-Trap beads (ChromoTek) overnight, followed by one wash in high-salt buffer (25 mM Tris, pH 7.4, 1% NP-40, 300 mM NaCl) and two washes in 25 mM Tris, pH 7.4, 1% NP-40, 150 mM NaCl. Proteins were eluted by in 1x Laemlli buffer (Bio-Rad) and heated to 95°C for 5 min. The samples were loaded on a 4-20% SDS Tris-Glycine gradient gels (Bio-Rad). When samples entered the gel by ∼1 cm, the run was halted, and the gel was fixed in ethanol-acetic acid-water solution (3:1:6 v/v). The gel plug was excised and submitted to the Keck MS & Proteomics Resource at Yale School of Medicine for LC MS/MS mass spectrometric analysis to identify interacting partners.

Briefly, excised SDS-PAGE gel corresponding to co-immunoprecipitated proteins was fixed with methanol/water/Acetonitrile (MeOH/H_2_O/CAN) (45/45/10) and washed with water and 100 mM ammonium bicarbonate in 50/50 ACN/H_2_O solution. Gel plugs were then treated with 4.5 mM DTT (20 minutes at 37°C, then allowed to cool to room temperature) and alkylated with iodoacetamide (for 20 minutes at room temperature in the dark). They were then washed with (a) 100 mM ammonium bicarbonate in 50/50 ACN/H_2_O solution, (b) 25 mM ammonium bicarbonate in 50/50 ACN/H_2_O solution, then with (c) 100% acetonitrile solution prior to complete drying using the SpeedVac. Trypsin was then added to the dried gel plug for enzymatic digestion in situ at 37°C overnight.

Tryptic peptides were then extracted with 500 µL 0.1% trifluoroacetic acid in 80/20 ACN/H_2_O for 15 min; solutions were transferred to a new tube and dried in a SpeedVac. These digests were reconstituted in Buffer A, and the resulting supernatant was injected onto a Waters NanoACQUITY UPLC coupled to a Q-Exactive Plus mass spectrometer containing a Waters Symmetry® C18 180 µm x 20 mm trap column and a 1.7 µm, 75 µm x 250 mm nanoAcquity™ UPLC™ column (35°C) for peptide separation. Peptide-trapping is performed at 5 µl/min, 99% Buffer A (100% water, 0.1% formic acid) and peptide separation is done at 300 nL/min with Buffer A and Buffer B (100% CH_3_CN, 0.075% formic acid). High-energy Collisional Dissociation (HCD) MS/MS spectra filtered by dynamic exclusion was acquired for the MS and MS/MS Q-Exactive Plus data collection. Orbitrap was used to detect all MS (Profile) and MS/MS (centroid) peaks. Resulting data dependent LC MS/MS data were analyzed using Proteome Discoverer 2.4 (ThermoFisher Scientific) for peak picking; and with Mascot search engine (v. 2.7 Matrix Science LLC.) for database search with results imported into Scaffold (v. 4.0, Proteome Software) for further visual inspection and analyses. Two or more unique peptides per protein were considered positive hits with a Mascot Score greater than 95% confidence level. The mass spectrometry proteomics data have been deposited to the ProteomeXchange Consortium via the PRIDE (Perez-Riverol et al., 2021) partner repository with the dataset identifier PXD041630.

### Animals

All mouse protocols including generation of talin R1R2 mutant knock-in mice and all experimental procedures were approved by the Yale University Institutional Animal Care & Use Committee.

### Generation of mutant mice

The talin 1 L638D/V722R R1R2 mutant (R1R2 mutant) mice were generated stepwise via CRISPR-Cas9 methods (*51–54*). Potential Cas9 target guide (protospacer) sequences in the vicinity of the L638 and V722 codons were screened using the online tool CRISPOR http://crispor.tefor.net (*55*) and candidates selected. Templates for sgRNA synthesis were generated by PCR from a pX330 template (Addgene), sgRNAs were transcribed in vitro and purified (MEGAshortscript, MEGAClear; ThermoFisher). sgRNA/Cas9 RNPs were complexed and tested for activity by zygote electroporation, incubation of embryos to blastocyst stage, and genotype scoring of indels at the target sites. The sgRNAs that demonstrated the highest activity were selected. Guide RNA (gRNA) sequences are as follows: 5’ guide: GCCCCGTCAGAACCTGCTGC and 3’ guide ACTAGCTGTTCTTGGCACAC. Based on these choices, a 1.6kb long single-stranded DNA (lssDNA) recombination template incorporating the L638D and V722R changes was synthesized (IDT). The sgRNA/Cas9 RNP + lssDNA was microinjected into the pronuclei of C57Bl/6J zygotes (*53*). Embryos were transferred to the oviducts of pseudopregnant CD-1 foster females using standard techniques. Genotype screening of tissue biopsies from founder pups used PCR amplification and Sanger sequencing to identify the desired base changes. In the first round, only the L638D change was incorporated, thus, founder mice with that change were bred and their offspring were used as embryo donors for a second round of targeting, in which only the 3’ sgRNA+Cas9 and a 148-base oligonucleotide were electroporated (*54*) to generate the L638D/V722R mutant allele. Germline transmission of the correctly targeted allele (i.e., both changes in cis) was confirmed by breeding and sequence analysis.

### Atomic force microscopy (AFM)

Mechanical properties in aortas from P24 mice were determined using AFM as described (*56*). A patch (≈5 mm^2^) of tissue from the ascending or descending aorta was attached to a coverslip, with the adventitial surface facing upward. All experiments were performed in HEPES-buffered solution, pH 7.4. Measurements were obtained on a Bruker MultiMode 8 AFM in contact mode using a cantilever (PFQNM-LCA-CAL, Bruker) (*57, 58*) with a tip radius of 70 nm and a nominal spring constant of 0.1 N/m. Force-deflection curves were obtained using Nanoscope III analysis Software for each aorta at 2 different areas per aorta. The data obtained were fit using a Hertzian model (*59*) to obtain a compressive elastic modulus under small indentations.

### Passive biomechanics and burst pressure tests

Standard biaxial biomechanical testing was performed as described (*60*). Briefly, the ascending thoracic aorta (from the root to the brachiocephalic artery) and the descending thoracic aorta (from first to fifth pairs of intercostal arteries) were isolated from P24 mice, cleaned of perivascular tissue, and their branches ligated with a 7-0 nylon suture to allow *ex vivo* pressurization. The aortas were mounted on glass cannulas and placed in a custom computer-controlled biaxial testing system, with temperature and pH maintained at physiological levels. Following four preconditioning cycles of pressurization and depressurization from 10 to 140 mmHg at the energetically preferred *in vivo* axial stretch, four cyclic pressure-diameter tests and three cyclic axial force-length tests were performed. The unloading portion of the last cycle of all seven pressure-diameter and axial force-length tests were fit simultaneously with a microstructurally motivated four-fiber family nonlinear stored energy function, as described (*61*). Values of mean biaxial wall stress and material stiffness were computed from the stored energy function and calculated at a common transmural pressure of 70 mmHg (∼80-85 mmHg equivalent *in vivo* luminal pressure). Next, the samples were re-cannulated on a custom blunt-ended double-needle assembly, stretched to the estimated *in vivo* axial length, and pressurized until failure to determine burst pressure (*62*), an indicator of wall strength.

### Histology

Aortas were fixed in 4% paraformaldehyde overnight, transferred to ethanol, embedded in paraffin and 10 µm sections cut. Sections were deparaffinized in xylene followed by washes in 95%-70% ethanol gradient. For picrosirius red (Abcam, ab150681) and Movat pentachrome staining (Abcam, ab245884), slides were stained as per manufacturer’s instructions. For immunostaining for collagen III, antigen was retrieved by heating to 95°C in antigen retrieval buffer (Vector Labs, H-3300), followed by permeabilization in 0.025% triton-X in PBS and blocking overnight in 10% donkey serum and 1% BSA. Sections were incubated overnight at 4°C with collagen III primary antibody (rabbit; ab7778; Abcam; 1:100), rinsed in TBS, pH 7.4, incubated with HRP-conjugated anti-rabbit IgG (goat; ab97080; Abcam; 1:500) for 1 h at room temperature and then rinsed. Slides were developed using 3,3’Diaminobenzidine (DAB) (Abcam; ab64238).

### AlphaFold structural modelling

To produce the structural model of mouse R2 in complex with ARPC5L, ColabFold (*63*) was used to run the 3D protein structure prediction tool, AlphaFold (*64*). The R2 and ARPC5L sequences were submitted to the software, structural models visualized in PyMOL and compared against the crystal structure of mouse R1R2 (Protein Data Bank accession no. 1SJ8 (*27*) and cryoelectron microscopy structure of the ARP2/3 1B5CL complex (Protein Data Bank accession no, 6YW6 (*65*). All figures were made using PyMOL (Version 2.5.2; Schrödinger, LLC).

### Statistical analysis

Statistical analysis was performed using GraphPad Prism version 8 (Graph-Pad Software Inc.) or Origin (OriginLab Corp.) with statistical significance denoted by NS for not statistically significant where p > 0.05, * for p ≤ 0.05, **for p ≤ 0.01, *** for p ≤ 0.001, and **** for p ≤ 0.0001. Data were analyzed by unpaired student’s t-test, Mann-Whitney test, or one-way analysis of variance (ANOVA) or Chi-square analysis as indicated in the figure legends.

## Acknowledgements

We thank members of the Yale Genome Editing Center for generating the talin R1R2 mutant mice. We also thank Dr. George Tellides and Qunhua Huang for help with mouse experiments. We thank the Keck MS & Proteomics Resource for providing the necessary mass spectrometers and the accompany biotechnology tools which is partly funded by Yale School of Medicine and by the Office of The Director, National Institutes of Health (S10OD02365101A1, S10OD019967, and S10OD018034). We also thank Yale pathology tissue services for help with paraffin-embedding, sectioning and staining of tissues.

## Author contributions

Investigation, methodology, data acquisition, validation, curation, formal analysis and visualization- M.V.L Chanduri and A. Kumar.

Data acquisition, validation, analysis and visualization- D. Weiss, N. Emuna, I. Barsukov, M. Shi and B.T. Goult.

Data acquisition and analysis- X. Wang, A. Datye and U.D. Schwarz

Resource generation and validation- K. Tanaka, S. Bai and T. Nottoli

Data acquisition and curation- J. Kanyo, F. Collin and T. Lam

Conceptualization - M.A. Schwartz

Project administration, formal analysis, supervision, funding acquisition- M. A. Schwartz, J.D. Humphrey.

Writing: original draft- M.V.L. Chanduri and M. A. Schwartz

Writing- reviewing and editing- M.V.L. Chanduri, A. Kumar, D. Weiss, X. Wang, A. Datye, T. Lam, T. Nottoli, B.T. Goult, J.D. Humphrey and M.A. Schwartz

## Data and materials availability

All data mentioned in the manuscript are available upon reasonable request.

