## Supplementary information for "Mechanosensing through talin 1 contributes to tissue mechanical homeostasis"

Manasa V.L. Chanduri. A. Kumar *et al*.

**This PDF file includes:**

Supplementary Text
Figs. S1 to S10

**Other Supplementary Materials for this manuscript include the following:**
Tables S1 and S2

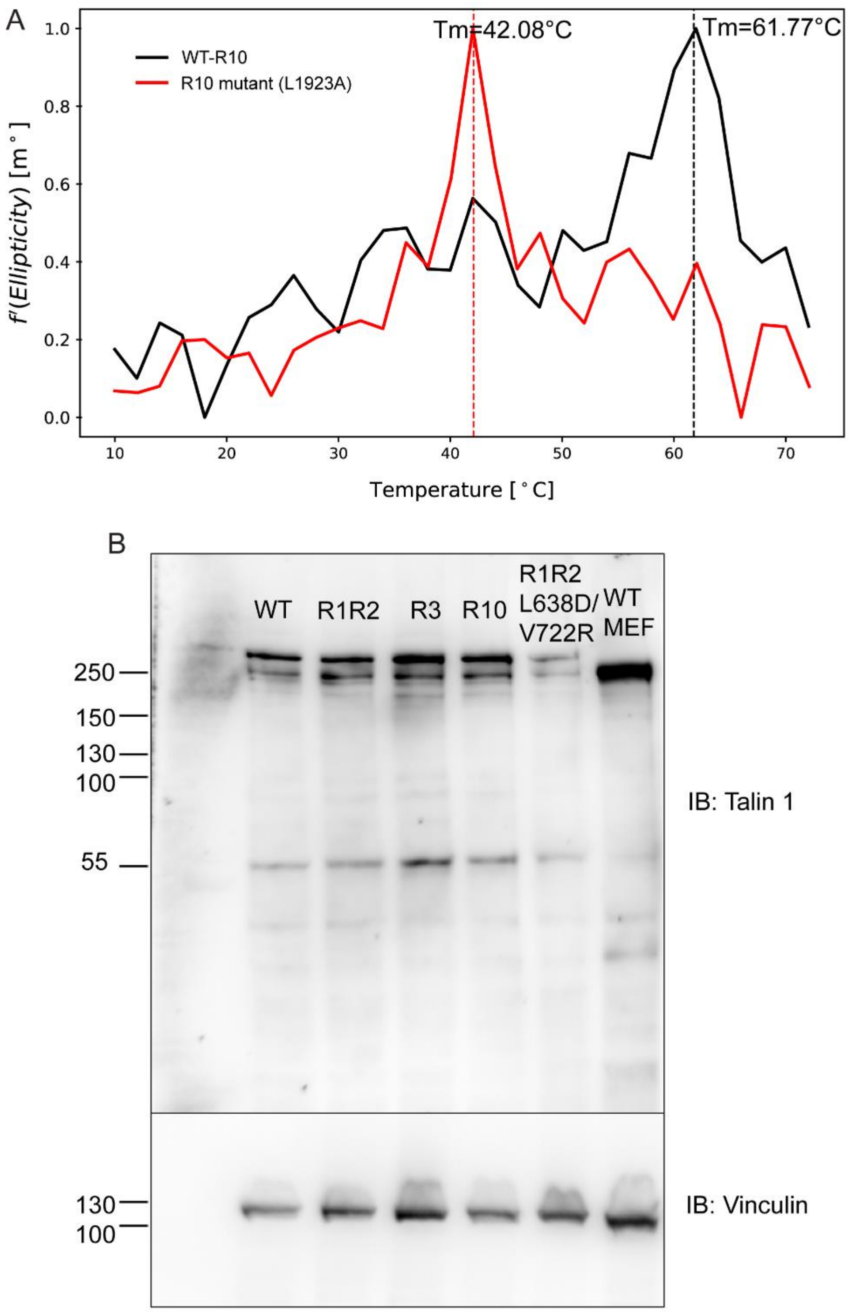


**Fig. S1. Characterization of R10 talin mutant**

A) CD spectra showing molar ellipticity at a wavelength of 220 nm vs temperature in WT and mutant R10 domain. Dotted lines indicate the melting temperature. B) Representative immunoblot for full-length WT and mutant talin containing the GFP-RFP FRET-sensor module {Kumar, 2016 #1} using anti- mouse talin antibody (mouse, ab104913, Abcam). WT MEFs were used as control for talin. Vinculin was used as a loading control.

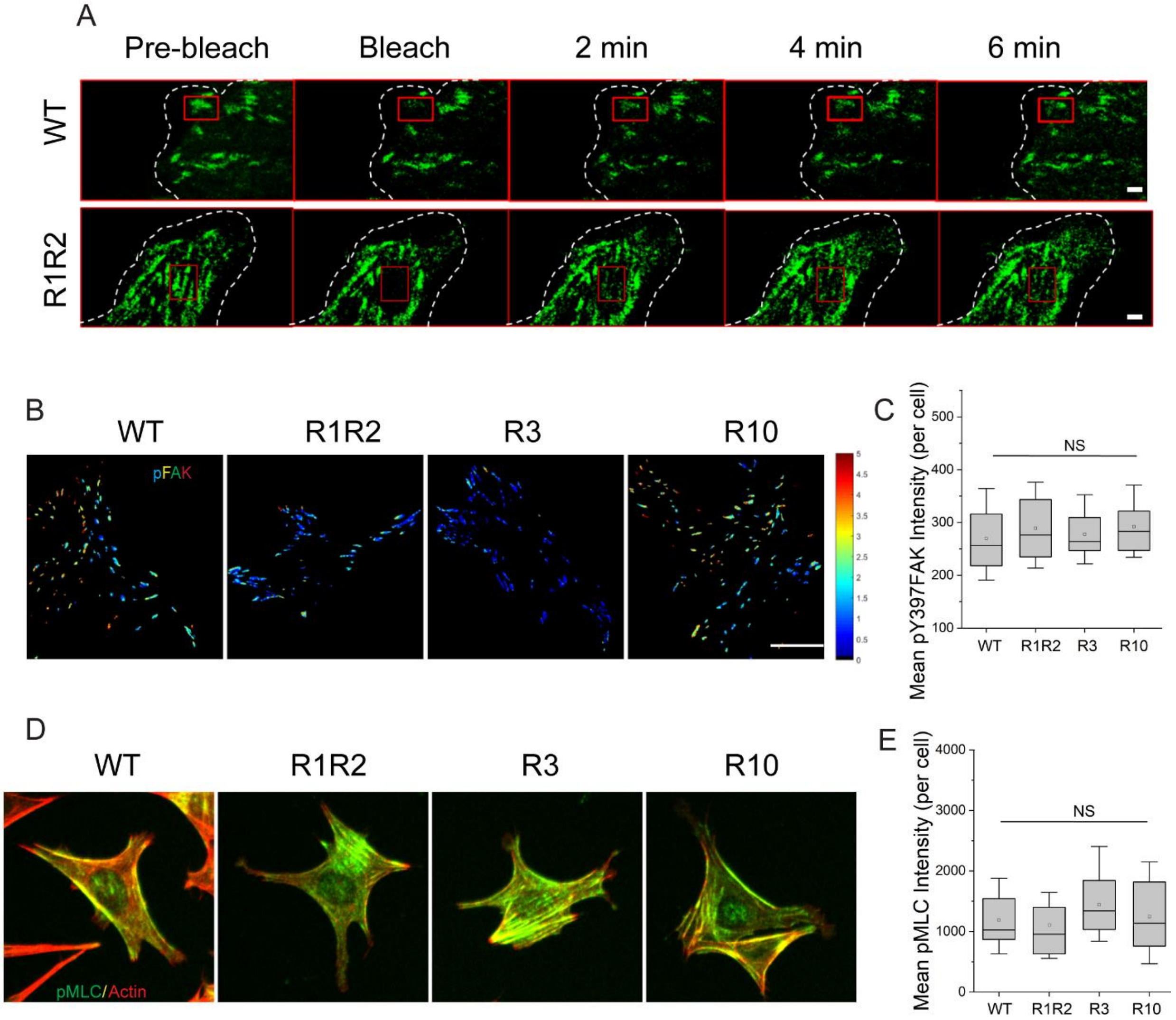


**Fig. S2: FA dynamics and signaling in talin rod-domain mutants.**

A) Representative images of FRAP experiment in *Tln1*$^{-/-}$ MEFs expressing full-length WT or the R1R2 mutant talin. Boxed regions indicate bleached regions. Scale bar, 1 μm. B) Representative ratio images of pFAK staining in *Tln1*$^{-/-}$ MEFs expressing full-length WT, R1R2 mutant, R3 mutant and R10 mutant talin. C) Quantitation of B). N=50 cells C) Representative immunofluorescence images of phospho-myosin light chain (pMLC; green) and phalloidin (actin; red) in *Tln1*$^{-/-}$ MEFs expressing full-length WT, R1R2, R3 and R10 mutant talin 1. D) Quantitation of C). N= 50 cells.

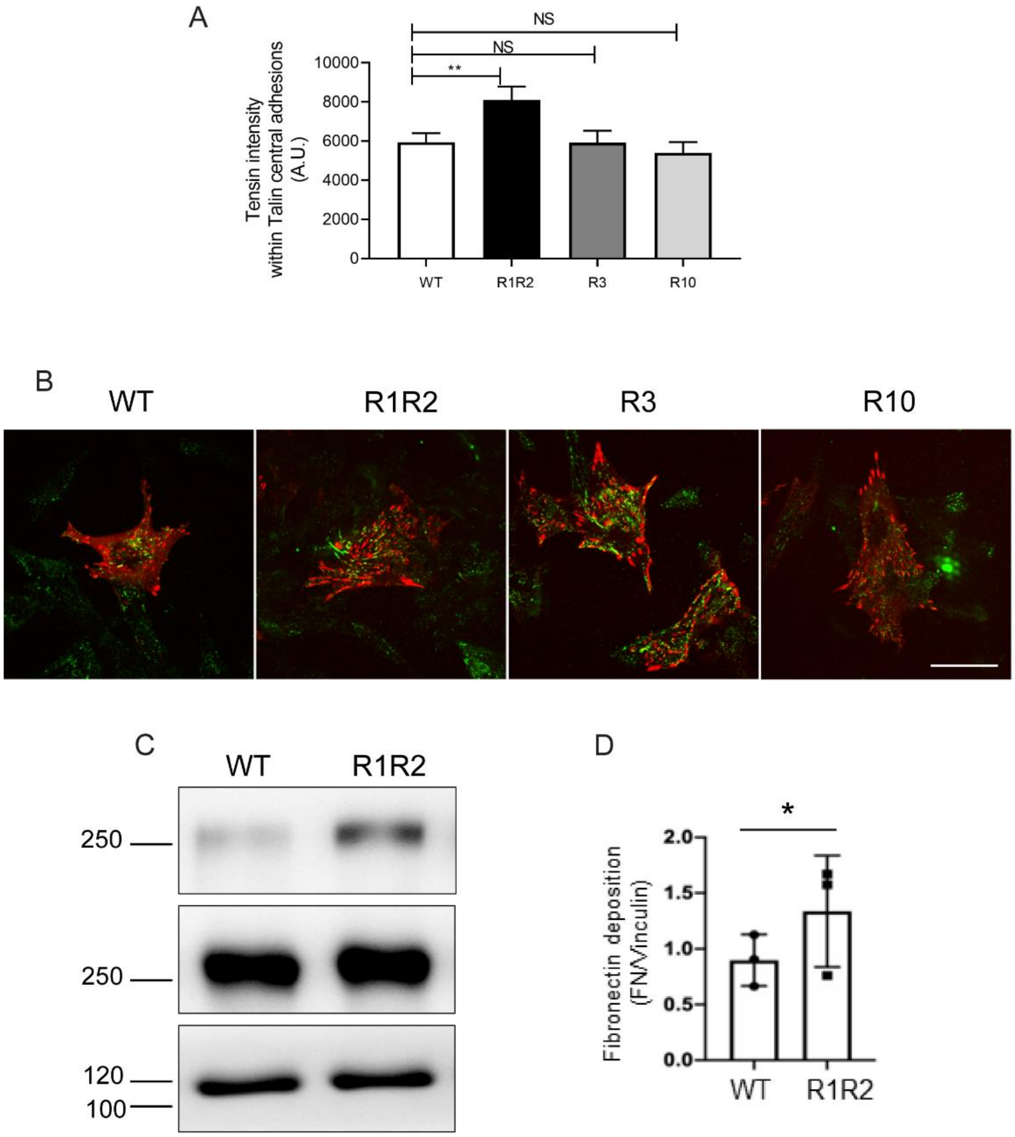


**Fig.S3: Fibronectin deposition in talin mutants.**

A) Tensin 1 intensity within talin-positive central adhesions. Values are means ± SEM of N= 25 cells. Statistics analyzed by one-way ANOVA. NS- Not significant; **, p<0.01. B) Representative images for figure 2F. Scale bar, 20 µm. C) Representative Western blot for detergent-resistant fibronectin in *Tln1*$^{-/-}$ MEFs expressing WT and R1R2 mutant talin. D) Quantification of C). N=3 experiments. Values are means ± SEM of N= 5. Statistics analyzed by unpaired t-test. NS- Not significant; *, p<0.05; **, p<0.01

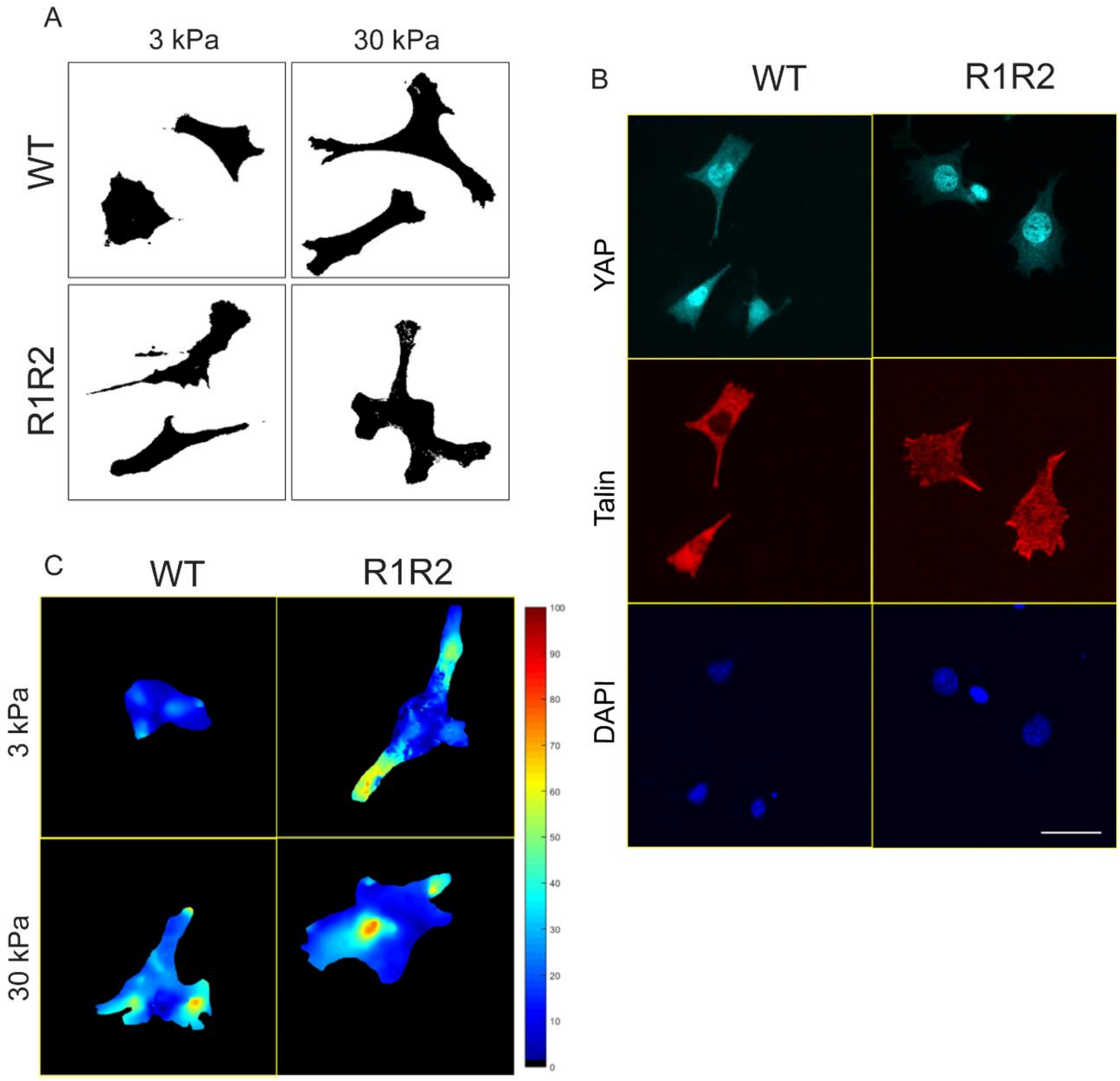


**Fig S4: Substrate stiffness-sensing in talin WT and R1R2 mutant cells.**

A) Representative inverted images of WT and R1R2 mutant cells stained with phalloidin on 3 and 30 kPa substrates. B) Representative images of YAP localization in WT and R1R2 mutant cells on 3 kPa. C) Representative traction force heat maps for WT and R1R2 mutant cells on 3 kPa and 30 kPa substrates.

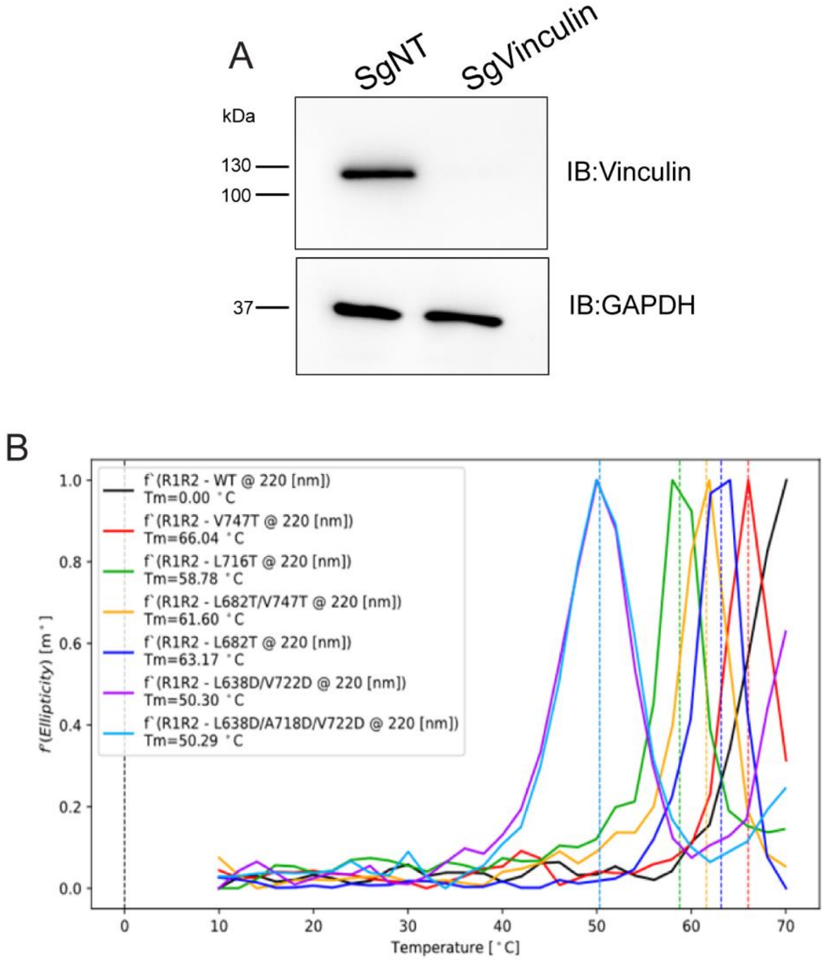


**Fig S5: Characterization of talin R1R2 core and interface mutations.**

A) Immunoblot showing knockout of vinculin in *Tln1*$^{-/-}$ MEFs. GAPDH (mouse, G8795, Sigma-Aldrich) was used a loading control. B) CD spectra showing molar ellipticity at 220 nm vs temperature in R1R2 core and R1R2 interface mutants compared to WT R1R2. Dotted lines indicate melting temperatures.

### Supplementary Figure 6

A

```
 ARPC5   MSKNTVSSARFRKVDVDEYDENKFVDEEDGG---DGQAGPDEGEVDSCLR
ARPC5L   MARNTLSS-RFRRVDIDEFDENKFVDEQEEAAAAAAEPGPDPSEVDGLLR
         *::**:** ***:**:**:********:: .    .:.*** .***. **

 ARPC5   QGNMTAALQAALKNPPINTKSQAVKDRAGSIVLKVLISFKANDIEKAVQS
ARPC5L   QGDMLRAFHAALRNSPVNTKNQAVKERAQGVVLKVLTNFKSSEIEQAVQS
         **:*  *::***:*.*:***.****:** .:***** .**:.:**:****

 ARPC5   LDKNGVDLLMKYIYKGFESPSDNSSAMLLQWHEKALAAGGVGSIVRVLTA
ARPC5L   LDRNGVDLLMKYIYKGFEKPTENSSAVLLQWHEKALAVGGLGSIIRVLTA
         **:***************.*::****:**********.**:***:*****

 ARPC5   RKTV
ARPC5L   RKTV
         ****
```

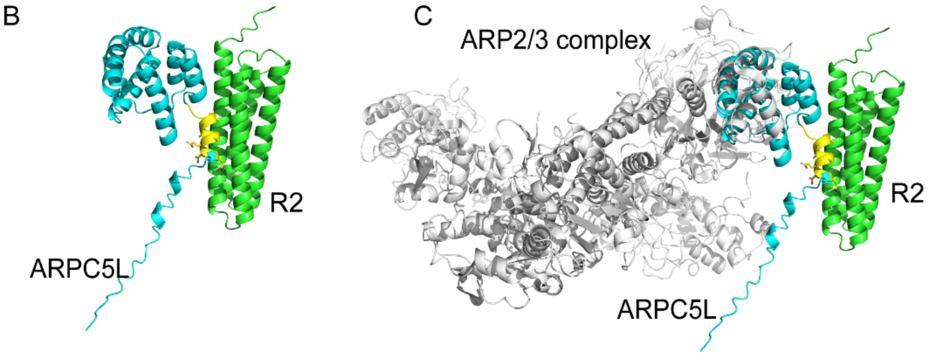


**Fig. S6: Modeling of ARPC5L interaction with talin R2 fragment.**

A) Alignment of ARPC5 and ARPC5L protein sequences. The region highlighted in yellow is unique to ARPC5L. B) AlphaFold model of the complex between talin R2 (green) and ARPC5L (cyan). The sequence highlighted in A) is shown in yellow. C) The talin R2-ARPC5L complex overlaid with the cryoelectron-microscopy structure of the ARP2/3 1B5CL complex (pdb 6WY6).

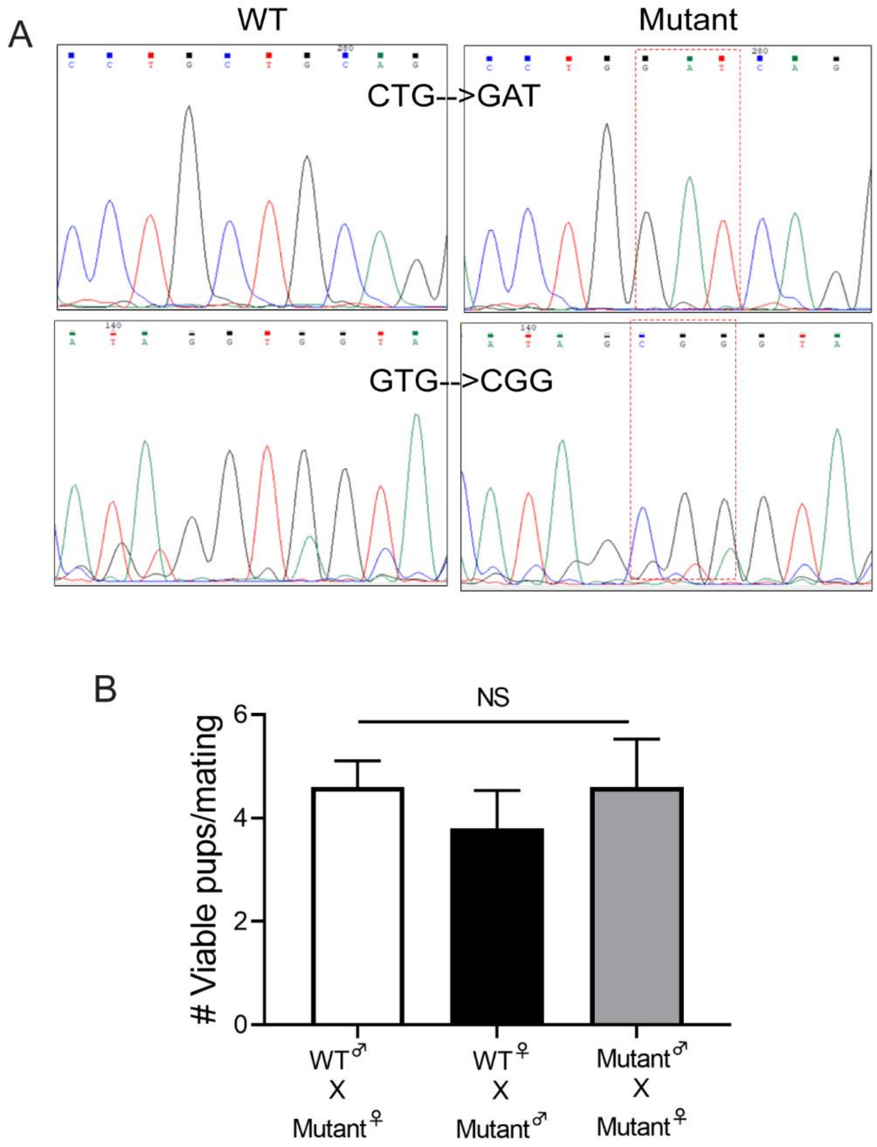


**Fig. S7: Characterization of WT and R1R2 mutant mice.**

A) Chromatograms of genotyped PCR products from WT and R1R2 mutant mouse showing the WT R1R2 and mutant R1R2 regions. Dotted lines indicate CTG to GAT (L638D) and GTG to CGG (V722R) knock-in mutations. B) Graph representing the viable pups from mating the R1R2 mutant male and female with each other and with WT male or female.

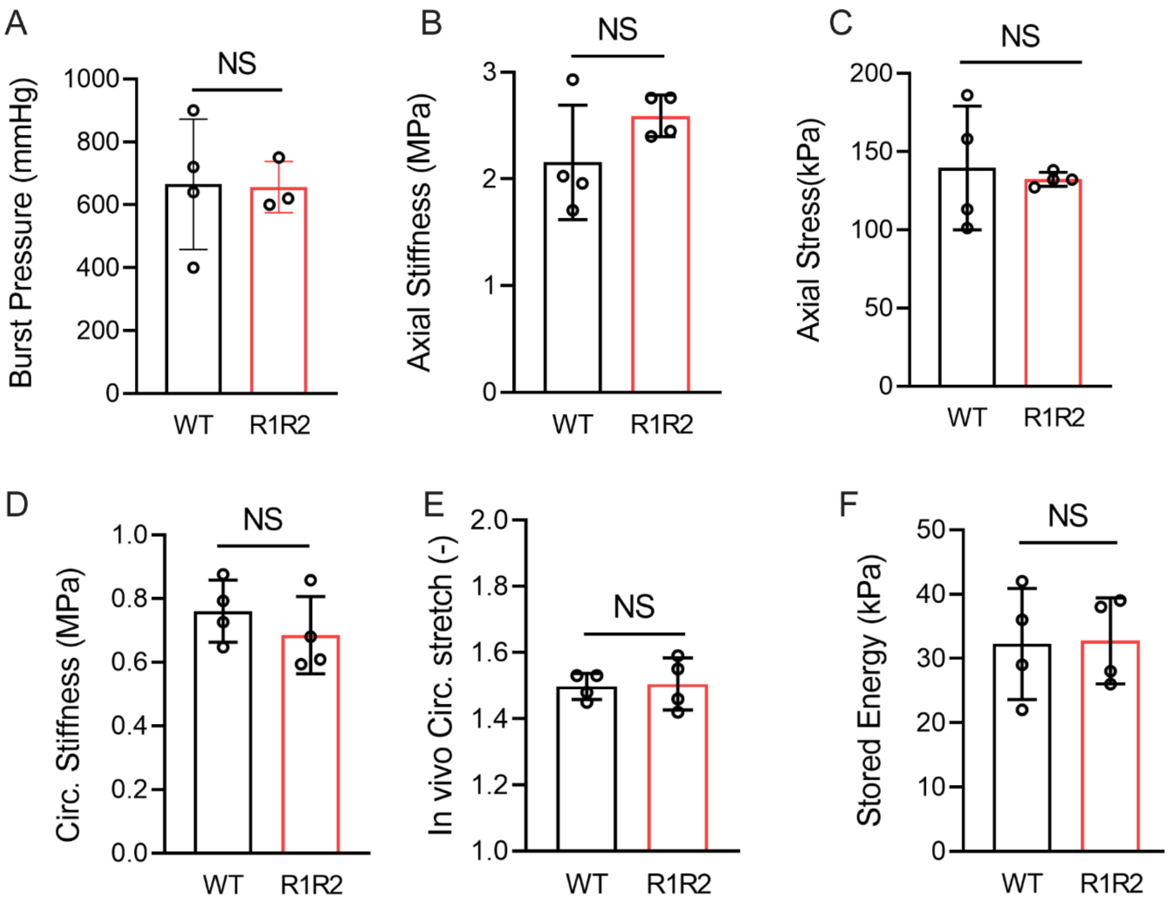


**Fig. S8: Mechanical analysis of descending aortas**

Biomechanical measurements of descending thoracic aortas from P24 WT and R1R2 mutant mice showing burst pressure (A), tensile axial stiffness (B), tensile axial stress (C), *in vivo* circumferential stretch (D), tensile circumferential stiffness (E) and stored energy density (F).

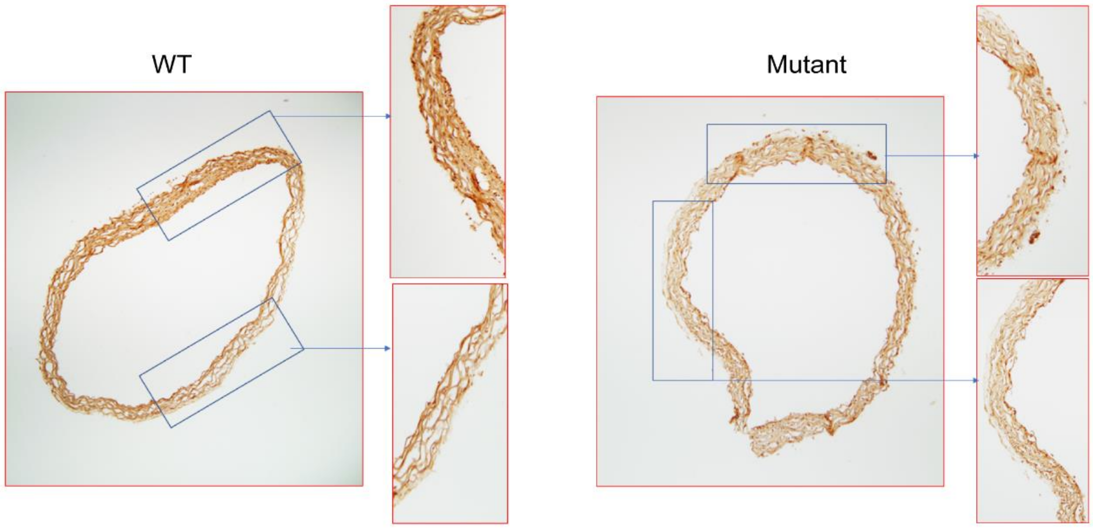


**Fig. S9: Collagen staining in R1R2 mutant mice.**

Representative images of collagen III immunostaining in WT and R1R2 mutant mouse ascending aorta cross-sections at P24.

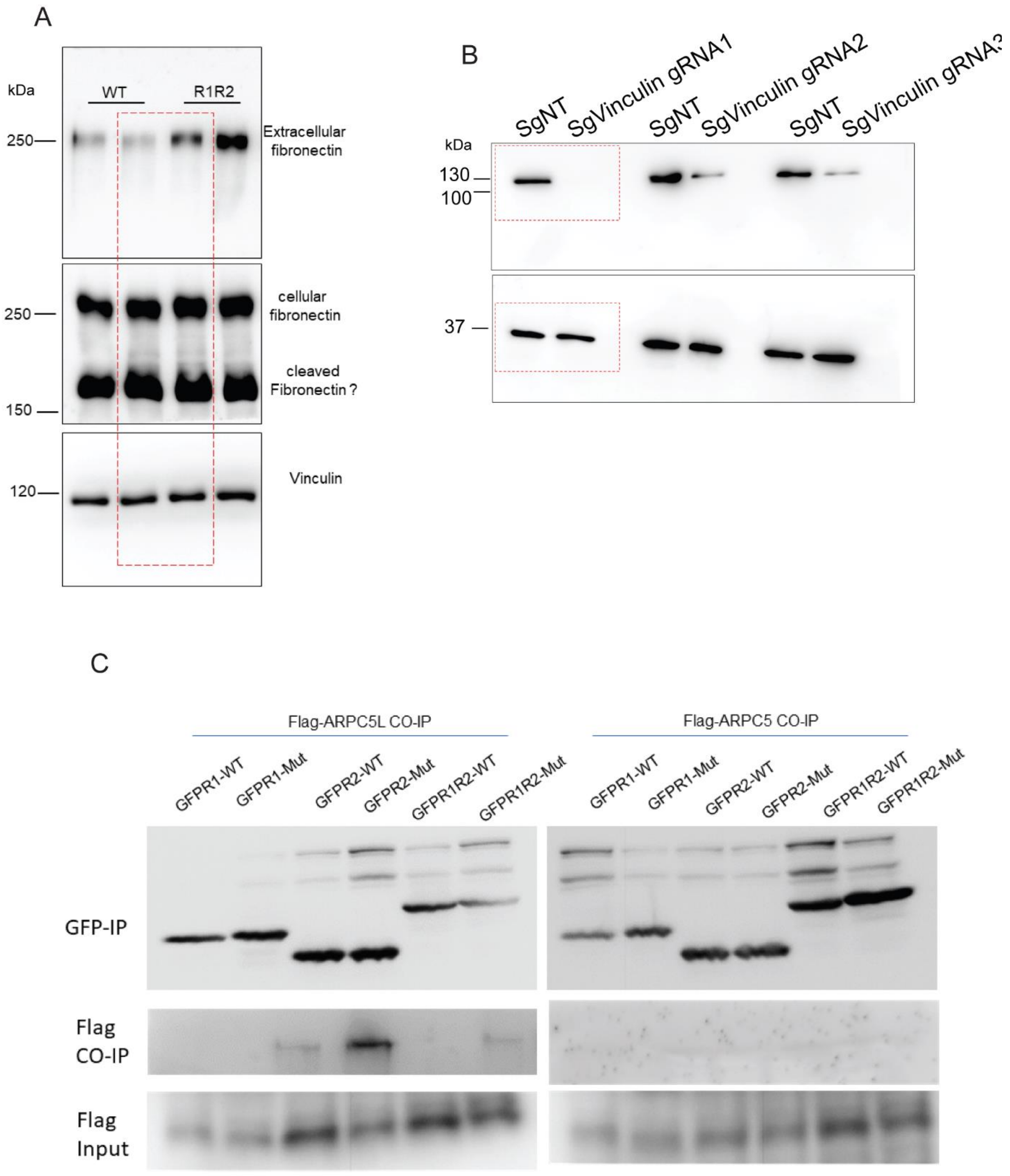


**Fig. S10: Full immunoblots:**

For figures S3C (A), S5A (B) and 6D (C).

**Supplementary tables (separate files):**

**Table S1:**
Complete list of proteins co-immunoprecipitated with GFP or GFP-tagged R1R2 or GFP-tagged R2 talin wild-type fragments based on unique peptide counts.

**Table S2:**
List of proteins that either exclusively bind or are enriched in GFP-talin R2 fragment samples.
